# Epileptic responses in marmosets induced by knockdown of CREB Regulated Transcription Coactivator 1 (CRTC1)

**DOI:** 10.1101/2023.09.22.558929

**Authors:** Yuki Nakagami, Misako Komatsu, Ken Nakae, Masanari Otsuka, Junichi Hata, Hiroaki Mizukami, Hiroshi Takemori, Shin Ishii, Hideyuki Okano, Akiya Watakabe, Tetsuo Yamamori

**Affiliations:** Laboratory of Molecular Analysis for Higher Brain Function, Center for Brain Science, RIKEN, Japan; Laboratory of Haptic Perception and Cognitive Physiology, Center for Brain Science, RIKEN, Japan; Brain Image Analysis Unit, Center for Brain Science, RIKEN, Wako, Japan; Integrated Systems Biology Laboratory Department of Systems Science, Graduate School of Informatics, Kyoto University; Kyoto, Japan; Division of Genetic Therapeutics, Center for Molecular Medicine, Jichi Medical University, Shimotsuke, Japan; Department of Chemistry and Biomolecular Science, Gifu University, Gifu, Japan; Department of Cellular Neuropathology, Brain Science Institute, Niigata University, Niigata, Japan; Central Institute of Experimental Animals, Kawasaki, Japan; Institute of Innovative Research, Tokyo Institute of Technology, Yokohama, Kanagawa, Japan; Exploratory Research Center on Life and Living Systems, National Institutes of Natural Sciences, Okazaki, Aichi, Japan; Faculty of Health Science, Tokyo Metropolitan University, Arakawa-ku, Tokyo, Japan; Keio University School of Medicine, Shinanomachi, Sinjuku-ku, Tokyo, Japan

**Author notes:** Corresponding author: Laboratory of Haptic Perception and Cognitive Physiology, Center for Brain Science, RIKEN, 2-1 Hirosawa, Wako, Saitama, 351-0198, Japan. These authors contributed equally (Yuki Nakagami, Misako Komatsu, Ken Nakae).

## Abstract

Animal models have contributed greatly to the development of anticonvulsant drugs. However, 20-30% of epilepsy patients still do not achieve adequate seizure control. It is important to understand the biological mechanisms of epilepsy in order to develop treatments and novel anticonvulsant drugs. Here we report *CRTC1* knockdown (KD) induced abnormal *cFOS* expression beyond the injection sites followed by epileptic response by three local injections of short hairpin RNA (shRNA) in marmoset V1. Longitudinal monitoring of cortical-wide neural activity revealed transient changes in high frequency oscillations by cortical electrocorticography throughout the cortex, and a lesion in the temporal lobe was observed by MRI and postmortem histology. *shCRTC1* KD marmosets showed severe lesions surrounding the injection site several months after injection. Glial cell activation may occur simultaneously or prior to IEG induction after *shCRTC1* injection. Thus, the epileptic process from seizure onset to remission can be studied in this animal model.

## Introduction

Epilepsy affects approximately 0.4-1% of the global population^1, 2^. Despite the invaluable contributions made by animal models combined with advances in drug discovery^3–9^, a subgroup of epilepsy patients, 20% to 30%, do not achieve seizure control or experience intolerable adverse effects^3, 8^. As most current epilepsy models are based on abnormal electrical potential changes or certain surface receptors, there is a growing demand to cultivate a variety of animal models, accurately representative of the 15 distinct epilepsy subtypes^9^ in order to facilitate the adoption of updated^10^ approaches in both research and drug development.

We have previously found immediate early gene (IEG) expression in the primary visual cortex (V1) of the marmose^11^ and simultaneous CRTC1 transient nuclear translocation and dephosphorylation of cAMP response element-binding protein (CREB) in V1 neurons following light stimulation (LS) in marmosets (unpublished data). Here, we report unexpected widespread *cFOS* expression in the injected hemisphere *CRTC1* KD induced by shRNA injections. A series of experiments ranging from histological analysis to cortical electrocorticography (ECoG) and magnetic resonance imaging (MRI) were performed.

The marmoset ECoG is advantageous for cortical-wide monitoring with high spatiotemporal resolution^12–14^ and long recording times. It allowed us to uncover sequential real time events after *shCRTC1* injection, and later histology showed that the cellular spread of neurodegeneration was localized to the injected area by activation of surrounding glial cells, while epileptic neuronal activity spread from the injection site throughout the cortex and abnormal functional connectivity was observed. Epileptic foci migrated to the temporal cortex, and functional connectivity eventually reached remission several months after injection. To our best knowledge, these features are the first to report the occurrence of epileptic focal transition, and make our shCRTC1 KD marmoset a strong candidate for a useful animal model of focal and structural epilepsy to study epileptogenesis in real time from onset.

## Results

This study started with the finding that *shCRTC1#1* KD induced unusual *cFOS* expression beyond the injection site. Our detailed methods are described in STAR METHODS. Briefly, we generated five shRNA sequences for KD of marmoset *CRTC1*, with *shCRTC1*#1 and #2 generated against the coding sequences and from #3 to #5 against the 3’ UTR sequences. *shCRTC1*#1 and #2 showed the strongest and second strongest *CRTC1* KD effects, respectively (Figure S1a). We used 10 marmosets in this study. The rearing conditions of these marmosets are described in the respective experiments and figures (see Supplemental Table 1a).

### *shCRTC1* injection at three local sites in V1 provides broad IEG expression

*shCRTC1*#1 together with Sirius was transduced by AAV1 into three sites in the left V1 (Figures 1a and 1b). A scrambled control sequence (*shscr)* was also transduced at one site and two sites in the right and left V1, respectively. Four weeks after shRNA injection, we first followed our previous condition^11^: That is, one eye was injected with tetrodotoxin (TTX) and subjected to dark rearing (DR) for 41hr followed by LS for 24 min (Figures. 1b). In contrast to *shscr*, *shCRTC1*#1 injection abolished CRTC1 protein expression (Figure 1c middle panel). In the control hemisphere with a single *shscr* injection, *cFOS* expression showed an ocular dominance column (ODC)-like pattern (Figures. 1c and d, bottom right and right, respectively). In contrast, aberrant *cFOS* expression was observed throughout V1 and V2 in the *shCRTC1*-injected hemisphere (Figures. 1c and 1d, bottom left and bottom left, respectively). *cFOS* mRNA expression in left V1 was strongest in layer 2, with the ODC pattern obscured by the abnormal cFOS expression and very different from the right hemisphere that showed the typical *cFOS* induction pattern observed under TTX and LS conditions^11^. These results demonstrated the widespread abnormal expression of *cFOS* in the *shCRTC1#1* injected visual cortex independent of the input to the visual cortex.

**Figure 1.**
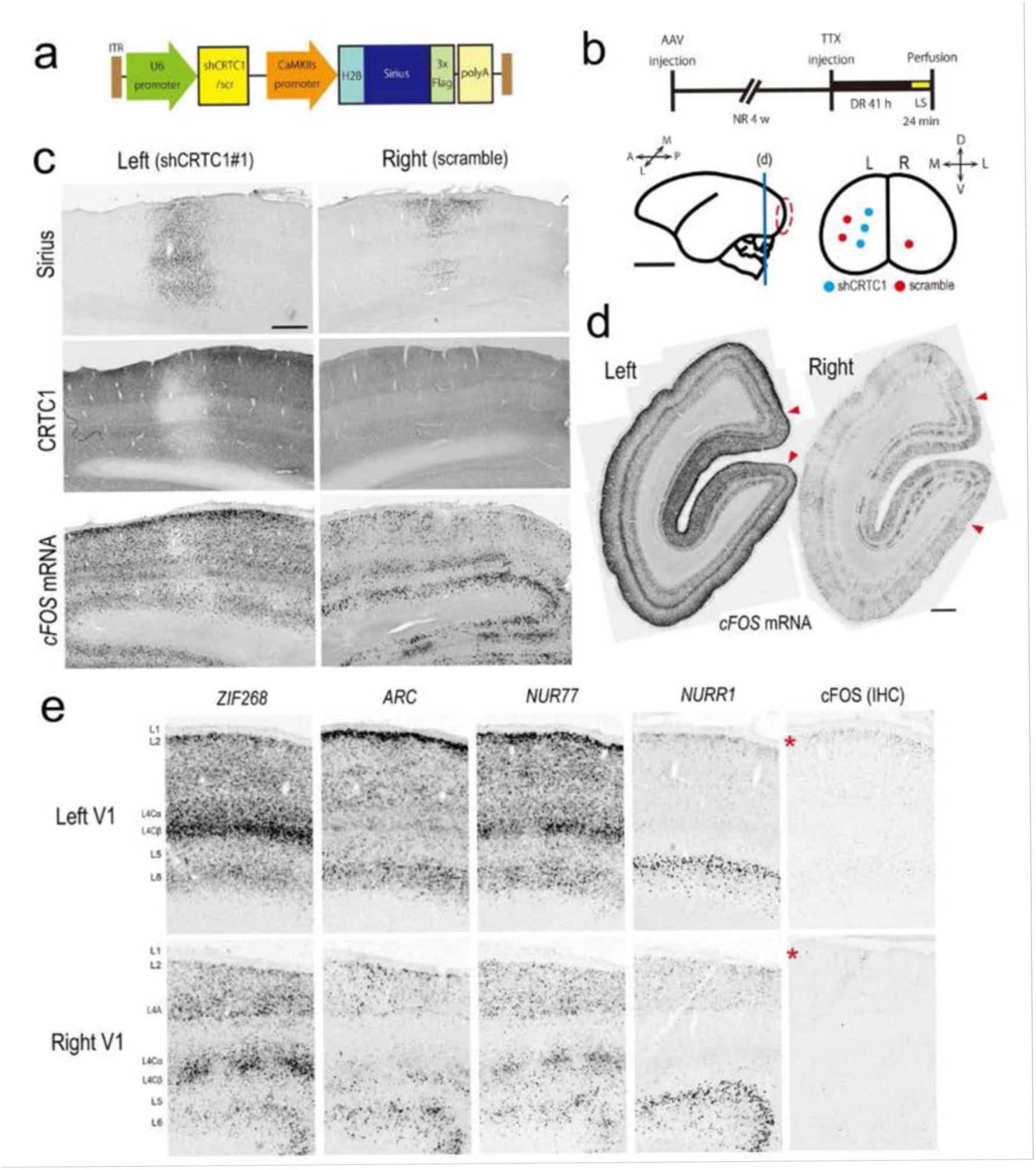
IEG induction in one hemisphere after injection of *shCRTC1*#l at three sites in Monkey 1. **a**, Construction *of the* AAV vector encoding *shCRTC1* (see STAR METHODS)*. **b**,* Upper: The experimental protocol (Monkey 1). Abbreviations: NR (normal rearing), DR (dark rearing), and LS (light stimulation). The time course is from left to right. Lower: Illustration of the *shCRTC1#1* injection sites. The *shRNA* was injected into the V1 region indicated by the red dashed circle from the lateral view. The right figure shows the injection site of for *shCRTCI* and *shscr* with red and blue filled circles, respectively. The injection sites in the marmoset V1 are surrounded by the red dotted line (left panel), shown in the posterior view (right panel). Abbreviations: A(anterior), P(posterior), L(lateral), M (medial), D (dorsal), and V (ventral). The blue line under the letters of ‘d’ marks the position of the coronal plane shown in d. Scale bar (1 cm). **c**, Expression of Sirius (IHC), CRTC1 (IHC), and *cFOS mRNA* (ISH) at the sh*CRTCI*#1 *(left)* and *shscr* (right) injection sites. Scale bar (0.5 mm). **d**, *cFOS* mRNA expression in coronal sections through the left and right V1. Red arrowheads indicate V1/V2 borders. Scale bar: 1 mm. **e**, Magnified views of the expression of five IEGs in the left and right V1. Sections adjacent to those examined for *cFOS* mRNA expression in Figure 1d were analyzed. cFOS protein was detected by IHC. Far right signals are for cFOS from ISH. Red asterisks indicate layer 2.

We examined the expression of several other IEGs at a distant site from the injection site within V1 (Figure 1e). The expression patterns of *ARC*, *ZIF268* (*EGR1* or *NGFIA*), and *NUR77 (NR4A1)* showed almost the same characteristics as those seen in the *cFOS* expression pattern (Figure S2). The strong expression in layer 2 with little ODC borders were different from those seen in the right V1. *NURR1* (*NR4A2*), a subtype of layer 6 neurons^15^ and the CREB downstream gene, showed activity-dependent gene expression in layers besides layer 6 at approximately 2hr after LS (data not shown). In the left V1 of Monkey 1, *NURR1* was aberrantly expressed only in the upper layer 2, without any abnormal expression pattern in other layers when compared to those of *shCRTC1* uninjected previous study^11^. In addition, the expression patterns of other retinal activity-dependent genes, *HTR1B* and *HTR2A*^11^, showed ODC-like patterns independent of *shCRTC1*#1 injection (data not shown), indicating that *shCRTC1*#1 injection induced more enhanced IEG expression in V1 independent of retinal activity.

LS-induced cFOS protein in the right V1 was undetectable at 24 min after LS, but significant cFOS protein expression was observed in layer 2 in the left V1 (Figure 1e, right), being consistent with the unusually enhanced IEG expression in the left hemisphere independent of LS. Around the *shCRTC1*#1 injection site, *ARC* and *ZIF268* (data not shown) were widely and uniformly expressed in addition to *cFOS* (Figure 1c, bottom left). *NUR77* expression was markedly reduced except in layer 2 at the *shCRTC1*#1 injection sites (Figure S1c), and the *NURR1* expression was slightly reduced in layer 6a and strongly reduced in layer 6b (Figure S1d). These results suggest that local reduction of CRTC1 altered local IEG expressions, which further affected much wider areas of the same hemisphere by the three-site injection of *shCRTC1*#1 in Monkey 1.

### Occurrence of abnormal high-frequency activity in *shCRTC1* KD marmosets

We consider that the wider expression of IEG may be accompanied with abnormal cortical activation. The abnormal IEG expression in layer 2 of Monkey 1 was similar to the IEG pattern reported in human tissue from epilepsy patients. In addition, the histological observations described below, such as the decrease in NeuN-positive cells, strong positive GFAP signals, and neuronal loss one month after three *shCRTC1*#1 injections (Figure S3b), were also consistent with those seen in human epilepsy^16^. We thus hypothesized that local injection of *shCRTC1*#1 into V1 induces abnormal excitatory activity in a cluster of infected neurons, which in turn causes epileptic activity that spreads throughout the cortex. To test this hypothesis, long-term recordings of cortical neuronal activity were performed in two *shCRTC1* KD marmosets (Monkeys 2 and 3). After injection of *shCRTC1*#1 at three adjacent sites in V1, ECoG electrodes^17^ were placed over the lateral hemisphere on the dura mater of Monkeys 2 and 3 on the same day and one week after injection, respectively (Figures 2a and 2b). After recovery, Monkeys 2 and 3 underwent ECoG recordings for 1-2hr per day and behavioral observations for 2 and 5 months, respectively. During the recordings, the monkeys sat in a primate chair without performing any task and were exposed to a variety of pure tones used in the previous study^13^ for the first 20 min. In addition to electrophysiological monitoring, preoperative *in vivo* MRI (T2-weighted imaging) and postmortem *ex vivo* MRI (T2-weighted imaging and diffusion MRI (dMRI)) were performed to examine structural changes (Figures 2a and 2b). Cortical activity was observed for 2 months in Monkey 2 and up to 5 months after the start of observation in Monkey 3. For quantitative analysis of the ECoG signal, we focused on the high-frequency (80-200 Hz) range obtained during the first 15 minutes of each day’s recording, because high-frequency oscillations (HFOs) are considered a biomarker for epileptic foci^16, 18, 19^. Examination of the high-frequency activity (HFA) (80-200 Hz, >3 SD) from all electrodes revealed that it occurred in multiple locations in V1, at the injection site, and even in regions of uninfected sites, and decreased 5 weeks after injection in both *shCRTC1* KD marmosets (Figures 2c-e). We then defined HFA spreading over more than 50% of the electrodes as cortical-wide HFO (cwHFO) (Figures. 2c, 2d and Figures S4 and S5). To identify the cwHFO onset site, we calculated the average value of the HFA 250ms immediately preceding the cwHFO. Early after *shCRTC1*#1 injection, HFA was observed in V1 preceding cwHFO, but by 4 weeks after the injection, these HFAs had decreased, and the HFA in the temporal cortex tended to precede the cwHFO (Figures. 2c, d and f, Figure S6 and Supplemental Videos 1 and 2). In addition, behavioral monitoring suggests that these HFAs are associated with behavioral outcomes. Both marmosets tended to look around and vocalize during periods of strong occipital HFA occurred but were stationary during periods of strong temporal lobe HFA occurred (visual observations). In addition, buccal hair ruffling was observed approximately two months after injection (visual observation).

**Figure 2.**
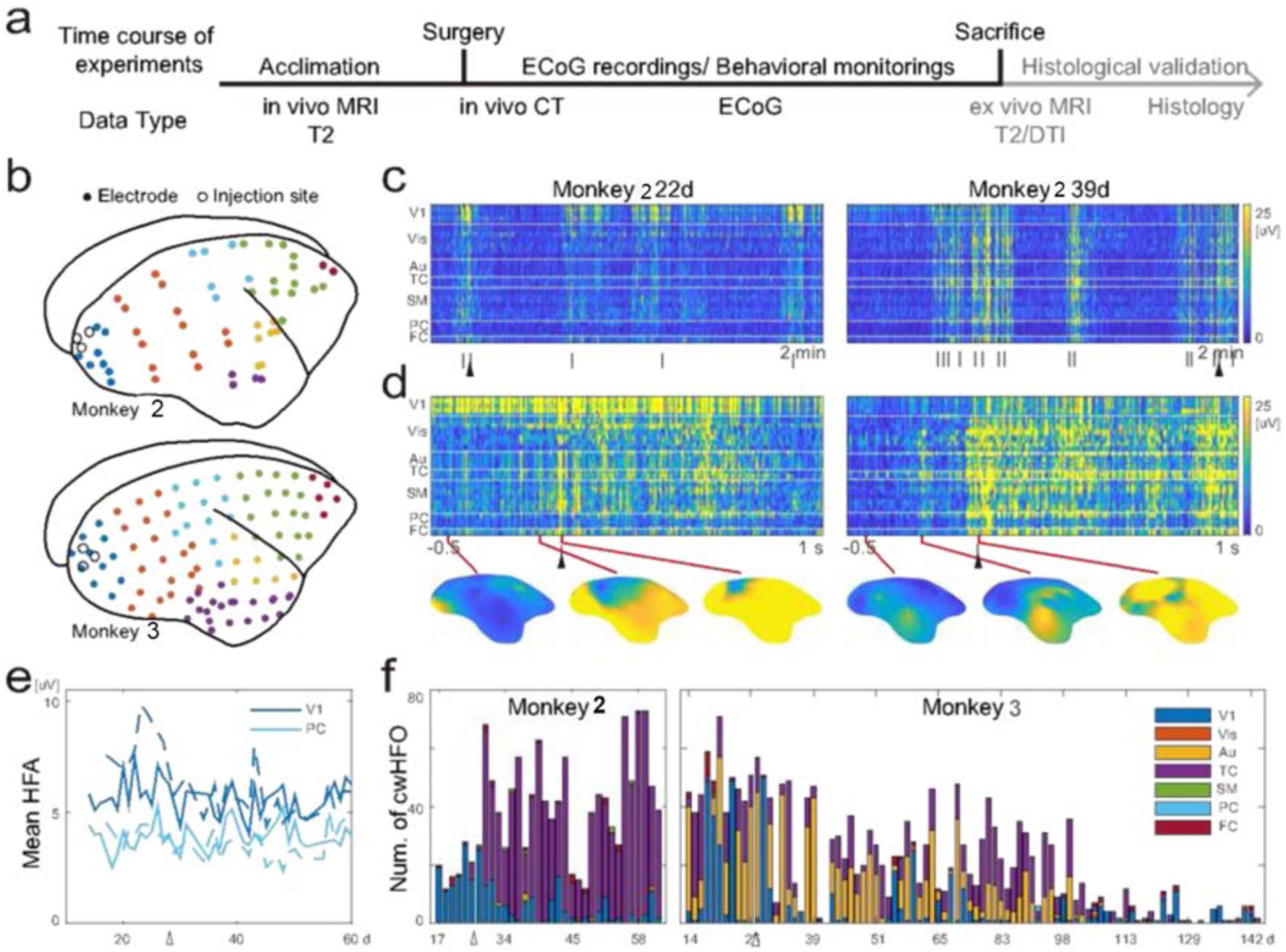
Local *CRTC 1*KD induced both local and cortex-wide HFOs. **a**, Time course of the experiments. **b**, Locations of the *shCRTC1*#1 injection sites (open circles) and ECoG electrodes (colored dots) in Monkeys 2 and 3. The colors of the dots indicate the grouped cortical regions: V1, Vis, Aud, TC, SM, PC, and FC. The color code is the same as shown in the inset of (f). **c**, Examples of HFA (80-200 Hz) in Monkey 2. The vertical bars at the bottom of the figures indicate the onset of cwHFOs. The filled triangles correspond to the cwHFO onset magnified in d. **d**, Examples of cwHFOs followed by HFOs in V1 (left) and TC (right). **e**, Mean HFA during 15 min recordings in V1 (blue) and PC (cyan) for each experimental day. The open triangle below the horizontal axis indicates 28 days after *shCRTC1* knockdown. **f**, Number of cwHFOs for each experimental day. The color of each bar indicates the cortical region that showed the highest HFA from -250 to 0ms before the onset of cwHFO.

### Reorganization of functional networks in epilepsy

To understand the significance of the changes in HFA patterns, we examined the daily functional network by calculating the correlations between each electrode on each recording day and averaging the correlations over all days (Figure 3a). The early recordings (22d and 18d) tended to show dissociations of the functional connectivity, or correlations, from the averaged functional connectivity after injection. In contrast, later recordings (56d and 55d) showed that the functional connectivity was close to the average (Figure 3b). To further investigate this variability, we compared the daily values of the average functional connection between presumptively defined cortical regions (see Methods). We found that the functional connectivity within 5 weeks after injection tended to differ from the average (Figures 3c and 3d). Among the connections, the value of the V1-V1 (other regions within V1) connection was significantly reduced in both monkeys (Figure 3e). This suggests that the local functional connectivity in the visual cortex decreased immediately after *shCRTC1*#1 injection and reorganized to a stable level five weeks after injection.

**Figure 3.**
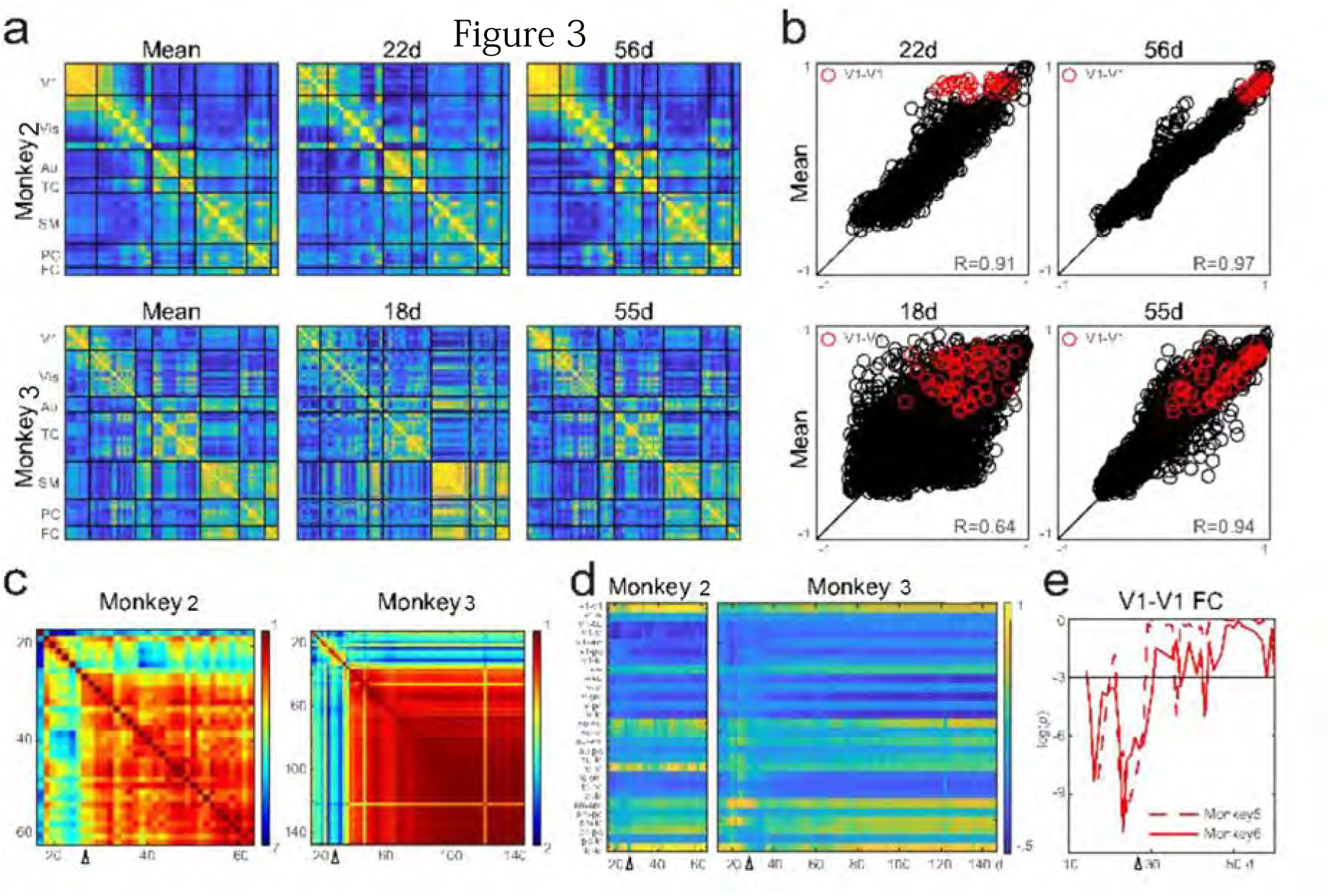
Altered functional networks reorganized 5 weeks after KD with shCRTC1#1. **a,** *CRTC1* KD induced alterations in functional networks. The left columns show the means of the correlation matrices of ECoG signals for Monkeys 2 (top) and 3 (bottom). Middle and right columns are the correlation matrices obtained three (22 days) and eight (56 days) weeks after KD, respectively. The color scale is the same as in d. **b**, Deviation from the averaged functional connection. Each functional connection is plotted on the x-axis for each experimental day and on the y-axis for the averaged network in a. **c**, Stability of functional connections. The left and right figures are the correlation matrices of the functional connections for each experimental day for Monkeys 2 and 3, respectively. The open triangles on the horizontal axis indicate 28 days after *CRTC1* knockdown. **d**, Functional connections between and within regions. **e,** Transient reductions in functional connectivity (FC) within V1. The horizontal line indicates the one-sided 5% significance level.

### *Ex vivo* dMRI reveals cortical asymmetry associated with reorganization of connectivity in monkeys injected with *shCRTC1*

We analyzed the *ex vivo* dMRI scans of Monkeys 2 and 3, obtained dMRI connectivity matrices among the whole-cortical supervoxels, and calculated the correlations between the connectivity profiles of the left and right hemispheres as asymmetric for 31 normal marmosets and Monkeys 2 and 3. We found that the *ex vivo* dMRIs of Monkeys 2 and 3 showed increased lateral asymmetry in connectivity, whereas the *ex vivo* dMRIs of the normal marmosets and the *in vivo* dMRI of Monkey 3 tended to show symmetry in most of the connectivity observed before the *shCRTC1* injection (Figure 4a). We then calculated z scores for the asymmetric features for 39 cortical regions of interest (ROIs). The ROIs that showed significant asymmetry in both Monkeys 2 and 3 were V1, V3, and S1 (Figure 4b).

**Figure 4.**
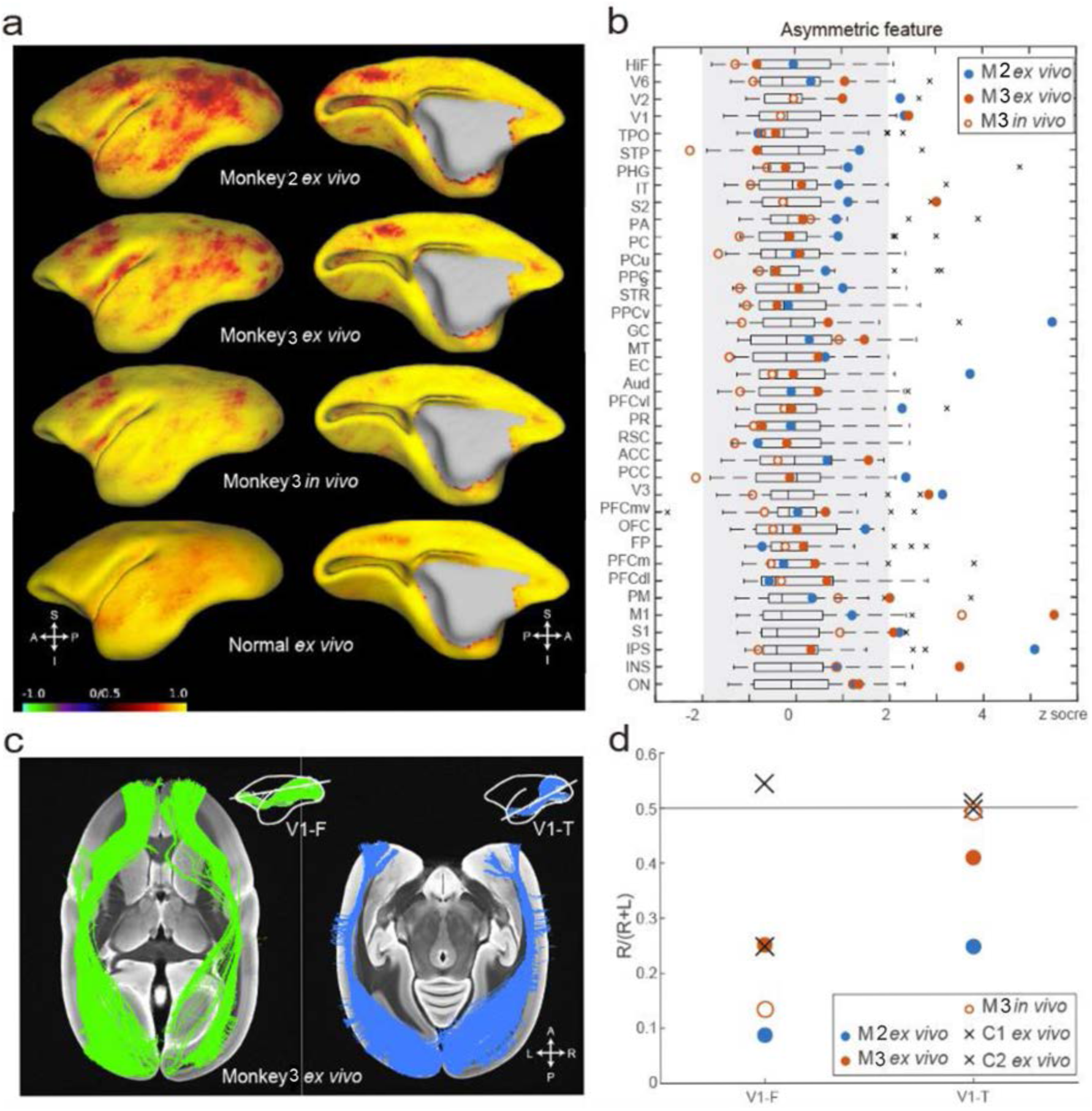
Postinjection dMRI shows cortical asymmetry. **a**, Correlations between the left and right profiles of dMRI connectivity at each supervoxel. The red region represents a low correlation, indicating asymmetric connectivity between the left and right cortices. **b**, The asymmetric feature (AF) values of 31 cortical ROIs. Box plot shows the AF from the *ex vivo* dMRI scans of 36 normal marmosets. The gray region represents a significant two-sided z score at a threshold of p<0.05 (-1.96 < z <1.96). Postinjection *ex vivo* AF of Monkey 2, the postinjection *ex vivo* AF of Monkey 3, and postinjection *in vivo* AF of Monkey 3 are indicated as filled blue, filled red, and open red circles, respectively. **c,** Diffusion tensor tractography. Fiber tracts connect V1 and the frontal (left) and temporal (right) areas of Monkey 3. **d**, Asymmetry of the fiber tracts. Abbreviations: HiF=hippocampal formation; V6=visual area 6; V2=visual area 2; V1=primary visual area; TPO=temporopolar area; STP=superior temporal polysensory cortex; PHG=parahippocampal gyrus; IT=inferior temporal area; S2=secondary sensory area; PA = area prostriate; PC=piriform cortex; PCu=precuneus; PPC=postal parietal area; S=subiculum; STR=superior temporal rostral area; PPCv=ventral postal parietal area; GC=gustatory cortex; MT=mid-temporal area; EC=entorhinal cortex; Aud=auditory cortex; PFCvl=ventral lateral prefrontal cortex; PR=perirhinal cortex; RSC=retrosplenial cortex; ACC=anterior cingulate cortex; PCC=posterior cingulate cortex; V3=visual area 3; PFCvm=ventromedial prefrontal cortex; OFC=orbitofrontal cortex; FP=frontal pole; PFCm=medial prefrontal cortex; PFCdl=dorsolateral prefrontal cortex; PM=premotor cortex; M1=primary motor area; S1=primary sensory area; IPS=intraparietal sulcus; INS=insular cortex; ON=olfactory nucleus; OB=olfactory bulb.

In addition, diffusion tensor tractography (DTT) analysis of white matter bundles in the V1-frontal (V1-F; Figure 4c left) and V1-temporal (V1-T; Figure 4c right) regions was performed to compare the number of extracted fiber tracts in the right and left hemispheres. A left-right imbalance of V1-T was observed only in the *ex vivo* MRI of Monkeys 2 and 3 and not in the *ex vivo* MRI of normal marmosets or in the *in vivo* MRI of Monkey 3. In contrast, a left-right imbalance of the V1-F bundle was observed in the *ex vivo* MRI of Monkeys 2 and 3 and even in the *ex vivo* MRI of a normal marmoset and the *in vivo* MRI of Monkey 3 (Figure 4d). Thus, we observed the structural changes in the whole cortical network of Monkeys 2 and 3 by compared to normal marmoset cortices.

### Progressive cell death at *shCRTC1*#1 injection sites associated with epileptogenesis

We performed postmortem histological analysis of Monkeys 2 and 3, two and five months after *shCRTC1*#1 injections, respectively, to examine histological changes occurred after long-term abnormal cortical activity. In both Monkeys 2 and 3, the expression pattern of *cFOS* mRNA showed almost the same distribution as in the NR condition, except that cells expressing relatively strong *cFOS* were sparsely observed in the upper layer 2 throughout the hemisphere (Figure 5a). Other IEGs examined (e.g., *ARC* and *BDNF*) were also similar (Figure 5b). At the injection sites, sparse but relatively strong *cFOS* expression was observed around the *shCRTC1#1* but not *shCRTC1*#2 injection site in Monkey 2 (see Figures 5a, 5c and 5f lower). No significant *cFOS* signal was observed at the *shCRTC1*#1 injection site in Monkey 3 (data not shown).

**Figure 5.**
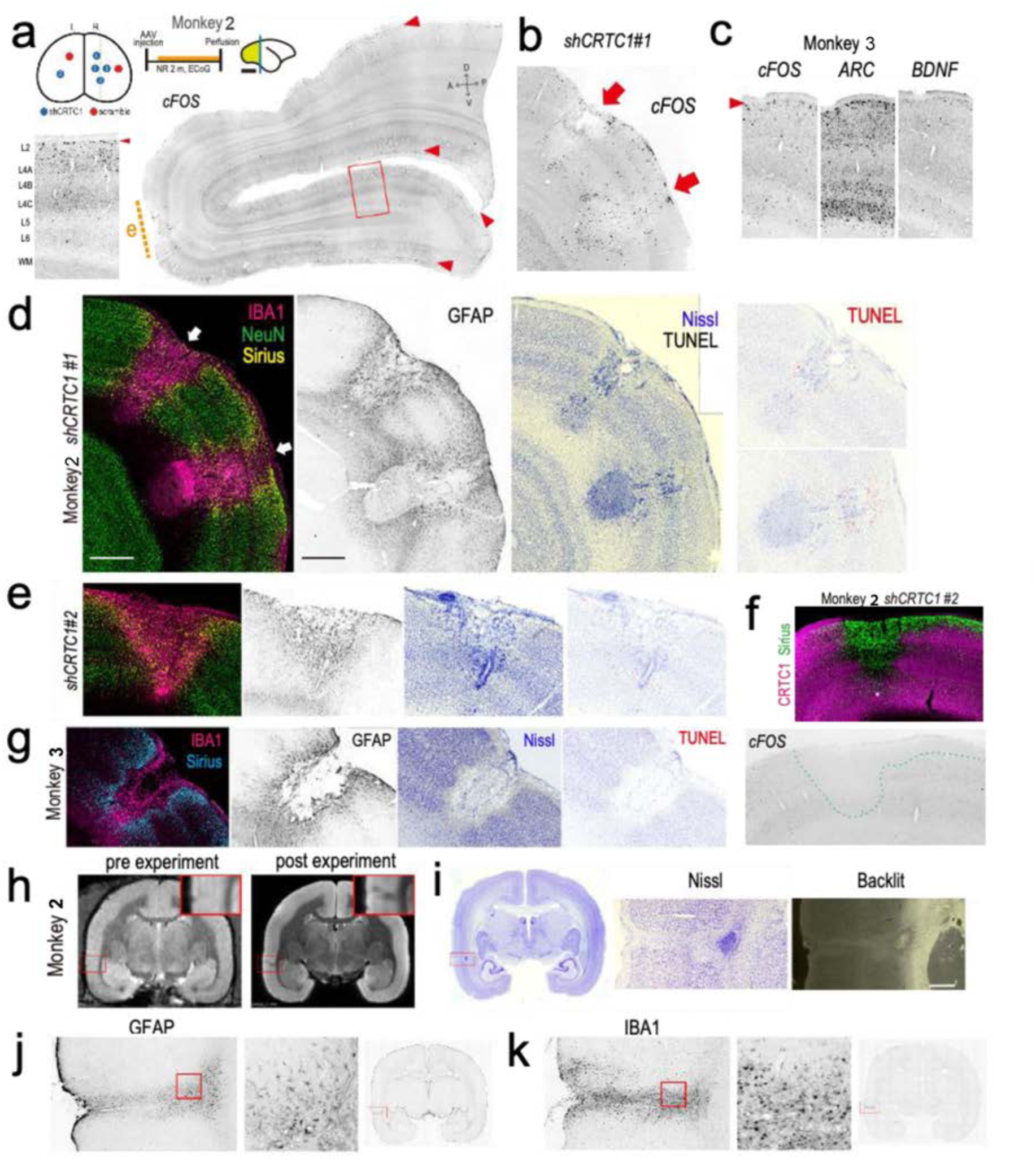
Histological analysis of 63 and 148 days after sh*CRTC1*#1 injection as recorded by ECoG. **a, c**FOS mRNA expression in a sagittal section of the right occipital lobe of Monkey 2. Upper left three small figures: Left (posterior view showing injection sites), injection sites of *shCRTC1*#1 (blue circles: three sites in the right hemisphere), *shCRTC1*#2 (blue circles: one site for each hemisphere) and *shscr* (red circles: one site for each hemisphere). Middle: Experimental time course. Right: schematic diagram showing the location of the section. Scale bar: 1 cm. The dotted green line (left; coronal view) and the yellow green area (right; sagittal view) indicate the position of the section. Bottom left: *cFOS* expression from the magnified view in the right section. Right: The large sagittal section shows the full view of the coronal section along with the blue line above. Red arrowheads indicate *cFOS* expression in layer 2. Orange dashed line “e” shows the position shown in “a”. **b,** Distribution of *cFOS* expression at injection sites of *shCRTC1#1* in Monkey 2. This section is the adjacent section of “d”. Red arrows indicate the *shCRTC1*#1 injection site, the same position as indicated by white arrows in “d”. **c,** IEG expression in V1 distant from the injection sites in monkey 3: left (*cFOS*), middle (*ARC*), right (*BDNF*). The red arrowhead indicates layer 2. **d-g,** The marmosets were histologically analyzed 63 and 148 days after *shCRTC1*#1 injections at three sites in the right hemisphere of Monkey 2 and Monkey 3, respectively (Supplemental Table 1a). (d) Monkey 2, injected with *shCRTC1*#1 at two injection sites (top left), was analyzed as shown in each of the four columns [IBA1, NeuN, and Sirius; GFAP only; Nissl and TUNEL; and TUNEL signals are displayed in red, from left to right]. Scale bar (500 μm). (e) Monkey 2, injected with *shCRTC1*#2 in the same staining as d. Adjacent sections with **a** were analyzed. (f) Expression pattern of *cFOS* at *shCRTC1*#2 injection site in Monkey 2. Upper: Merged image of double IHC of CRTC1 and Sirius signals. Bottom: ISH of *cFOS*. (g) Monkey 3 in the same staining as (d) without NeuN staining. **h**, *In vivo* (left) and *ex vivo* MRI (right) images of Monkey 2. The lesion in the temporal cortex (presumably TE3) is magnified in the upper right inset of each image. **i,** Images of Nissl staining with low magnification (left), high magnification (middle) and “backlit” light microscopy (right). **j,** GFAP signals. Left: the red boxed region of the left images in “i”. Middle: High magnification of the region indicated by the red box in the left image. Right: A lower magnification view of the section. **k,** IBA1 signals shown in the same order as the GFAP signals in “j”. Adjacent sections were analyzed in “i” and “j”.

In Monkey 2, *shCRTC1*#1 and *shCRTC1*#2 were injected (Figure 5d, and Figures 5e and 5f, respectively). Strong IBA1 signals and a mesh-like structure with GFAP signals were observed only at the edge of the injection site where Sirius-positive neurons were observed (Figure 5d). The strong IBA1 signal suggested undergoing phagocytosis of dying neurons (Figure 5d). GFAP signals forming reticular structures indicate activated astrocytes undergoing gliosis (Figure 5d). This suggests that cell death was well advanced before two months after injection, beyond the time point at which we examined histology. Monkey 2 was sacrificed five months after injection. Most of the neuronal loss was within the surrounding wall. GFAP signals also disappeared in the center of the injection, forming a glial scar^20, 21^ (Figure 5g). As shown by Nissl staining, neurons in most injection sites were completely lost, and TUNEL signals were not observed at the injection site or its periphery (Figure 5d, right).

Postmortem MRI T2 images of Monkey 2 showed signal loss in part of the right temporal lobe (Figure 5h), but no such signal loss in *in vivo* MRI T2 images taken before *shCRTC1*#1 injection; Nissl staining also revealed columnar-like cell death in the right temporal lobe of Monkey 4 (Figure 5i). We detected signals for GFAP and IBA1 (Figures. 5j and 5k, respectively), but no TUNEL-positive neurons (data not shown). These results suggest that the cell death events may peak about two months after the *shCRTC1*#1 injection and most cells die out in the injection center and possibly temporal cortex (Figures 5h and 5i).

### Comparison of IEG Induction and cell death markers for CRTC1 KD at one or two sites vs. injections at three sites per hemisphere

Considering that each injection site is approximately 1 mm in diameter and the marmoset cortex measures approximately 3 cm x 2 cm (M-L) (Figure 1a), AAV injection sites occupy only a small fraction of V1. We thus examined how the total number of injection sites affected IEG induction in Monkeys 1, 4, and 5, which were three sites, two sites and one site injections, respectively per V1 hemisphere (Figure 1, Figure S3, and Figures. 6a and 6b). As mentioned above, a population of *ARC* and *ZIF268* positive signals were uniformly and weakly expressed similar to *cFOS* (Figure S2). Strong signals of *cFOS, ARC*, *ZIF268* were observed in upper layer 2, but not at the *shsc*r injected site. *NURR1* showed in a way that it was attenuated only in layer 6b of the injection site (Figure S1d), but there was little other *shCRTC1*-induced *NURR1* layers besides layer 6 and upper layer 2 (Figure 1e, Figure S1d, and Figure S2). Monkey 4 normally reared 4 weeks after injected at two sites in the left hemisphere showed the strong *cFOS* induction at the injection site, especially in layers 2/3, overlapped with Sirius signals and weak cFOS expression beyond the injection site (Figure 6a). Monkey 5 underwent normal rearing for 4 weeks after AAV injection, followed by DR and LS before sacrifice (Figure 6b). LS induced broad *cFOS* mRNA expression, with much stronger intensity around the right V1 *shCRTC1*#1 injection site (Figure 6c, middle). In contrast, only limited *cFOS* and *ARC* inductions and *NUR77* reduction were observed with a much less injection size at the left V1 induction site (Figure 6d, middle). Note that *NUR77* was also strongly reduced at the injection site, but with induction outside the injection site in monkey 1 (Figure 1e and Figure S1c). To summarize the effect of *shCTRTC1*#1 injections on IEG expression, monkeys 4 and 5, although strong cFOS protein expression was observed (Figures 6a and 6c), the stronger *cFOS* induction than LS induced *cFOS* did not spread far beyond the injection site and did not appear to cause epileptic-like responses in monkeys 4 and 5.

**Figure 6.**
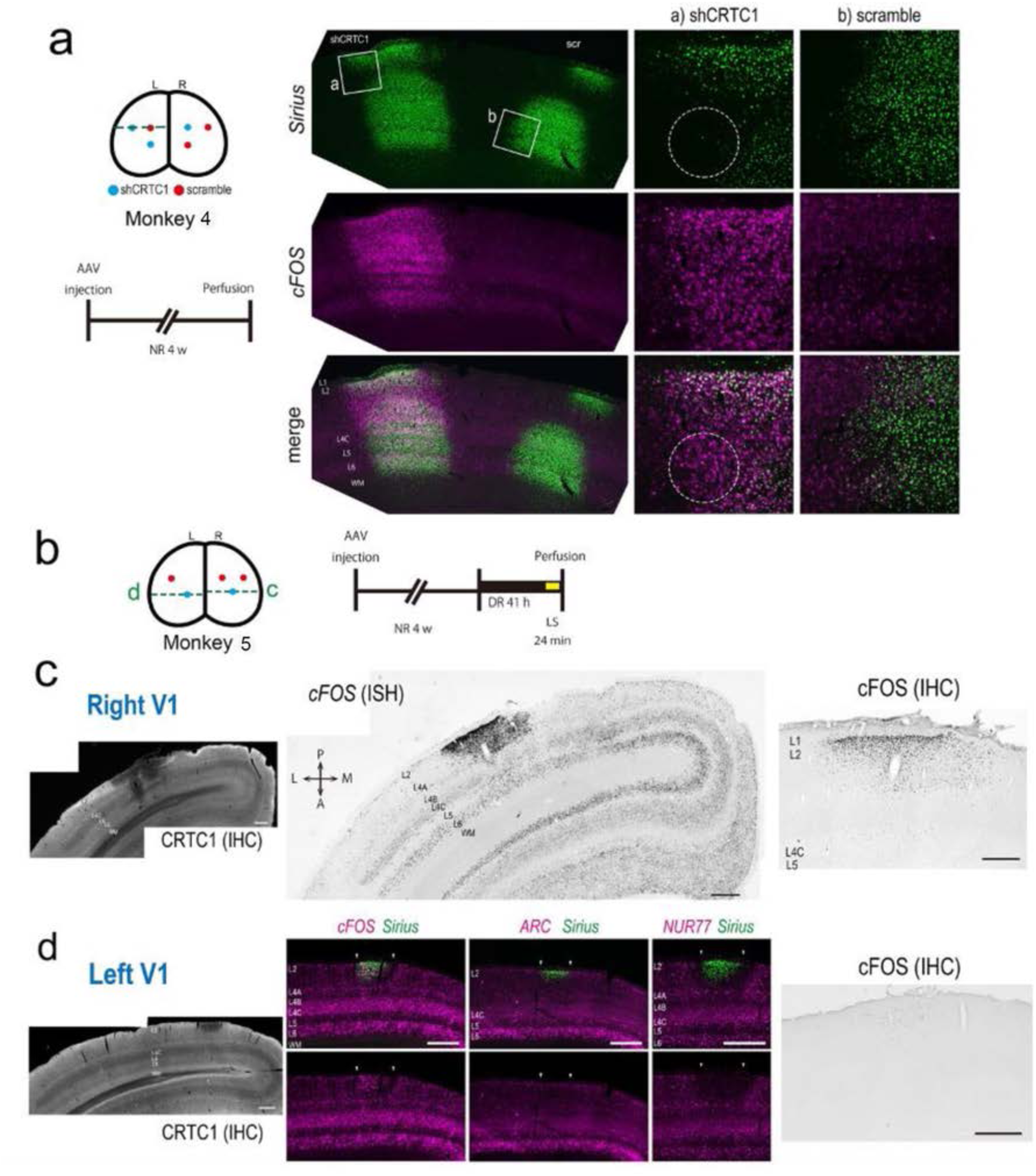
cFOS expression after injection of *shCRTC1*#1 into one site per hemisphere. **a,** Upper left panel: Three injections were made into each hemisphere; two of *shCRTC1*#1 and one of *shscr* into the left (L) hemisphere and one of *shCRTC1*#1 and two of *shscr* into the right (R) hemisphere of a normally reared animal (Monkey 4). Three right panels: Top row: Sirius (green) indicates the expression in the AAV-transduced cells. Middle row: *cFOS* (ISH) was detected at the *shCRTC1*#1 injection site but lower at the site of the *shscr* injection site. Bottom row: Merged images of *Sirius* and *cFOS* signals. **b,** Left: *shCRTC1*#1 was injected into one site in each hemisphere (Monkey 5), and *shscr* was injected into one site and two sites in the left and right hemispheres, respectively. Right: A normally reared marmoset that underwent 24 min of LS without a TTX eye injection (Monkey 5). **c-d,** Histological analyses of Monkey 5. (c) Left (right V1): CRTC1 expression detected by IHC. Middle: *cFOS* expression (by ISH). Strong and relatively weak *cFOS* expression was observed at the injection site and through the layers, in response to the *shCRTC1* injection and LS, respectively. Right: cFOS expression detected by IHC. Scale bars (200 μm). (d) (left V1): Left panel: CRTC1 (IHC). Middle panels (from left to right): *cFOS, ARC*, *NUR77* (magenta) with Sirius signals (green). Upper row: merged images. Lower row: IEG expressions. Right panel: cFOS (IHC). Scale bar (200 μm). L2-6 and wm (layers 1-6 and white matter, respectively). Red and blue filled circles in (a) and (b) indicate the injection sites for *shCRTC1*#1 and *shscr* RNAs, respectively. The other abbreviations are the same as in Figure 1b.

### Expression of cell lesion and apoptosis related markers 24 days after injection in Monkeys 1, 4 and 5

Because we detected severe neuronal loss conspicuous in Monkey 2 and 3, we examined the extent of neuronal lesions induced by *shCRTC1*#1 injection, together with other cell injury or death markers in Monkeys 1, 4, and 5. Nissl staining revealed microglial accumulation and tissue damage, and NeuN signaling was reduced and IBA1 and GFAP were increased at the *shCRTC1*#1 transduced site in Monkey 1 (Figures S3a and S3b), with little or no signal reduction seen with *shscr* injections (data not shown). The NeuN signal was attenuated in *shCRTC1*#1-trasduced cells and was markedly reduced in areas of severe tissue damage. Consistent with Nissl staining, IBA1 signal, indicating microglial activation, was observed at the center of the injection site. In contrast, the GFAP signal indicated astrocyte activation around and near the edge of the injection site (Figure S3a, right). Cell death signals detected by TUNEL staining, were in the central region of the *shCRTC1*#1 injection site in Monkey 4 (Figure S8b). No TUNEL signal was observed at the *shscr* injection site (data not shown). Because CRTC1 activation and CREB phosphorylation are independent processes induced by synaptic activity^22^, we examined the phosphorylation status of CREB at SER133 by IHC. Significant phosphorylated CREB signals were observed in neurons injected with *shCRTC1* (data not shown). We also performed a series of analyses on neurodegeneration in Monkeys 4 and 5. Nissl staining showed less neuronal loss and tissue damage in the two Monkeys than Monkey 1. Correspondingly, the immunostaining signals for IBA1 and GFAP were weaker than Monkey 1 (Figures S3a and S8a). NeuN signals were also reduced at *shCRTC1* injection sites, but only a few TUNEL signals were observed (Figures S3b and S3c, Figures S8b and S8c). pCREB signals were also observed at the s*hCRTC1* injection sites in Monkey 1 (Figure S3d), Monkeys 4 and 5 (data not shown) and Monkey 6 (shown later in Figure S9f). Overall, injection of *shCRTC1*#1 at one or two sites was weaker in IEG induction and cell death related marker expression than that of injection at three sites in V1.

### Expression of cell death related markers and IEG induction in each species of five different *shCRTC1*s in one site per hemisphere

To evaluate the effect of the other *shCRTC1*s besides *shCRTC1*#1, we injected different *shCRTC1*s and *shscr* per one V1 hemisphere of Monkey 6 (Supplemental Table 1a). Twenty-four days after *shCRTC1*s injection, Monkey 6 received a TTX injection in one eye, was maintained in DR for 2 days, and was light-stimulated for 24 minutes before sacrifice (Figure S9a), similar to Monkey 1. All *shCRTC1*s, but not *shscr*, reduced CRTC1 signals (Figure S9b). Cell death related and IEG expression markers were examined. Among variable NeuN reduction occurred at each injection site of different *shCRTC1*s (Figure S9c), more histological damage with *shCRTC1*#1 and #2 were observed than with other *shCRTC1*s (Figure S9d). Strong IBA1 signals were present at sites #1 and #2, with complementary GFAP signals (Figure S9e). *ShCRTC1*#1 and #2 also showed phosphorylated CREB signals (Figure S9f). *cFOS* mRNA expression decreased at all *shCRTC1*s injection sites (Figure S9g and data not shown), with no abnormally elevated IEG expressions. In contrast, we observed IEG induction at one injection site of *shCRTC1*#1 in Monkey 4. We interpret this difference that only half of the V1 areas were activated compared in Monkey 6 under NR and LS after 41hr of monocular TTX injection compared to normal marmosets. In the former condition, LS induced *cFOS* expression as an ODC-like pattern in V1, in which the reduction of *cFOS* at the *shCRTC1*#1 injected sites may be due to the reduced activated area of half of the *shCRTC1* injected region in V1 due to monocularly injected TTX in monkey 6. This may be consistent with the finding that less *shCRTC1*#1 injection resulted in less *cFOS* induction (Figures 6c and 6d). Monkey 7, normally reared, received *shCRTC1*#2 and #3 injections at one site each and shCRTC1#5 at two sites (Figure S9h). IEG expression was reduced at *shCRTC1* but not *shscr* injected sites (*ZIF268*, Figure S9h) without abnormal IEG expression. Monkey 8 confirmed this with *shCRTC1*#2 and #5 injections (Supplemental Table 1a). Extensive *shCRTC1*s-mediated IEG expression was also seen in Monkeys 9 injected with *shCRTC1*#1, along with *shCRTC1*#2 and #3, at one or two sites that alone cannot cause such IEG induction (Supplemental Table 1a). The difference between the effects of *shCRTC1*#1 and *shCRTC1*#2 seems to be due to that *shCRTC1*#1 influenced more IEG induction, while *shCRTC1*#2 may have influenced more tissue damage.

### *shCRTC1*#2 induced *cFOS* expression beyond the injection site when injected at 2 or 3 sites per hemisphere

To further characterize the effect of *shCRTC1*#2 compared to that of *shCRTC1*#1, we injected *shCRTC1*#2 at two or three sites per each V1 hemisphere in Monkey 10 (Figure 7a). We found *cFOS* expression beyond the injection site in both hemispheres, with increased IBA1 and decreased NeuN signals at the injection sites (Figure 7). Three-site injection of *shCRTC1*#2 further induced *cFOS* expression, decreased NeuN expression and increased IBA1 and TUNEL signals; however, compared to three-site injection of *shCRTC1*#1 in Monkey 1 (Figure 1d), *cFOS* expression was more restricted to the periphery of the *shCRTC1*#2 injection site (Figure 7b).

**Figure 7.**
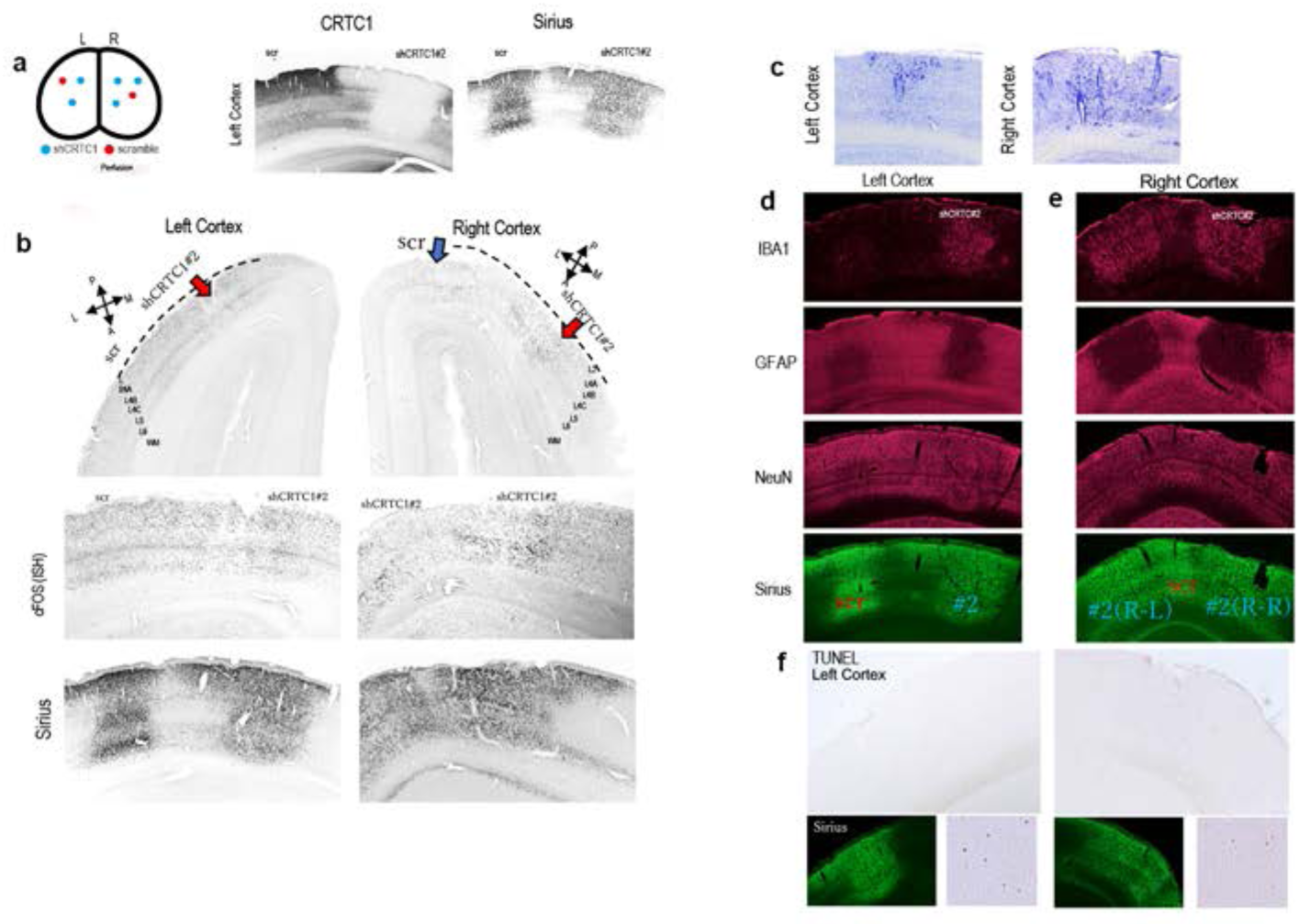
*cFOS* expression at *shCRTC1*#2 injection sites. *cFOS* expression at *shCRTC1*#2 injection sites. **a,** Left: the injection sites in V1 hemispheres (top) and in Monkey 10 (see Supplemental Table 1a for the rearing condition). Middle and right panels: IHC signals of CRTC1 and Sirius at the *shscr* and *shCRTC1*#2 injection sites. **b,** Top row: A low magnification view of cFOS expression as detected by ISH. L: lateral, M: medial, P: posterior, R: rostral. The *shCRTC1*#2 injection sites are indicated by red arrows, and the *shscr* injection site is indicated by a blue arrow (left hemisphere injection site indicated as scr). Note that *cFOS* expression was extended beyond the *shscr* injection site in the left cortex, and that the right V1 injections of *shCRTC1*#2 and *shscr* partially overlap in the cutaway view. Middle and bottom rows; Larger views of the V1 cortex below the dotted regions shown in the top figures are rotated approximately 90° clockwise (left cortex) or counterclockwise (right cortex). Middle and bottom rows show *cFOS* and Sirius signals detected by ISH and IHC, respectively, in the adjacent section. Scale bars: 500 μm. **c,** Nissl staining at the *shCRTC1* injection sites in the left and right cortices, respectively. **d,** Injection into left V1 cortex. Expression of IBA1, GFAP, NeuN, and Sirius at the injection site from top to bottom. In the bottom Sirius (IHC) staining figure, the injection sites of *shscr* and *shCRTC1*#2 are indicated by red and blue letters, respectively. **e,** Injection into the right V1 cortex. IBA1, GFAP, and NeuN signals are shown in magenta. Sirius signals are shown in green (bottom). Injection sites for *shscr* and *shCRTC1*#2 are shown using red and blue letters, respectively. Note that in the view of the bottom figure (Sirius), the injection sites for *shCRTC1*#2 and *shscr* are partially overlap due to the angle of the cross-section, as indicated by the purple arrow in a. **f**, Top row: Low-magnification TUNEL stained images of the left and right V1 cortices. Bottom row: Each image pair shows Sirius and enhanced TUNEL signals (in the region shown in the top image) in the left and right cortices.

The activity induced by injection of *shCRTC1*#1 was not strong when the size of the infected cells was limited but was increased to unusually strong levels with multiple injection sites. Other *shCRTC1*s than *shCRTC1*#1 cannot induce a widespread IEG expression beyond the injection site by each single site injection alone due to the lower KD efficiency. Nonetheless, *shCRTC1*#2 can induce the aberrant expression of IEGs when the transduced cells and sites were clustered (Figure 7b), with significant cell lesions (Figure 7c), IBA1 and GFAP expressions, NeuN reduction (Figures 7d and 7e), and TUNEL signals (Figure 7f). It should be also noted broader IEG expression was observed by injecting *shCRTC1*#1 at one site and with other *shCRTC1*s at a few nearby sites as described above (Monkeys 7 and 9, Supplemental Table 1a).

## Discussion

We report here that three-site injection of *shCRTC1*#1 into marmoset V1 caused epilepsy that spread throughout the cortex on the unihemisphere with the following time course: (1) Expression of cell death related markers and neural lesion around the injection site were observed with one injection per hemisphere for all *shCRTC1*s. (2) When *shCRTC1*#1 was injected at one site per hemisphere into a normally reared marmoset, cFOS was induced at and around the injection site. (3) IEG induction spread throughout V1/V2 of the injection site hemisphere when *shCRTC1*#1 was injected at three sites in V1. (4) Spread of electrical activity throughout the cortex, and (5) formation of lesions at the injection sites and the temporal lobe. We observed neuronal loss, activation of astrocytes and microglia by GFAP and IBA1, respectively, and lesions surrounded by the glial scar. (6) ECoG recordings over the cortex suggested that HFOs were induced at the *shCRTC1* injection site, spread around it, and eventually propagated throughout the cortex.

cwHFOs were first induced around the injection site, then gradually from the temporal cortex, and finally more from the temporal lobe. The association between HFAs and several behavioral outcomes shown in the results confirms that the abnormal neuronal activities induced by three sites of *shCRTC1#1* injections were indeed epileptic responses, as these associations were observed repeatedly over a month. In Monkey 2, MRI and Nissl staining confirmed tissue lesions in the temporal lobe, suggesting epilepsy focal to the temporal lobe (Figures. 5h and 5i). On the other hand, epilepsy symptoms in this model went into remission within a few months after injection. Five-month follow-up of Monkey 3, from 3 months onwards, the frequency of cwHFOs decreased significantly (Fig. 2f, and Figures S7a and S7b). However, sporadic local HFOs continued to be observed in large areas of the cortex, which were absent in normal marmoset (Figure S7c). This may provide new insights into the relationship between intercortical network reorganization and remission of epilepsy symptoms.

Structural changes long time, several months after, *shCRTC1* injection were evident not only in the histology of the temporal lobe of Monkey 2 (Figure 4d), but also in the e*x vivo* dMRI-based tractography of both *shCRTC1*#1-injected marmosets (Figure 4). Analysis of the correlation between the left-right profiles of the *ex vivo* dMRI connectivity showed lower values in *shCRTC1*#1-injected Monkeys 2 and 3 than in normal marmosets; the lateralization of the V1-F fiber tractography was confirmed by *ex vivo* MRI in normal marmosets and by *in vivo* MRI in Monkey 3. Despite these structural changes, we observed a reduction in cwHFO and stabilization of functional connectivity.

At later stages (2 and 5 months) of the *shCRTC1*#1 injection, activated astrocytes may act as an anti-inflammatory agent with glial scar formation by astroglia, preventing further tissue damage from spreading^23^. It has been reported that activated microglia that appear early after status epilepticus, are required for the induction of epileptogenic astrocytes that appear late after status epilepticus^24^. In our case, when the efficacy of the injected *shCRTC1* was limited by injection size, reduced efficiency of each *shCRTC1*, monocular TTX injection, or a combination of these conditions, activation of astrocytes and microglia could be appeared even before the induction of IEGs was detected. Furthermore, strong IBA1 and GFAP signals were observed at the *shCRTC1* injection site in all subjects. *shCRTC1* injection could eventually cause a situation similar to status epilepticus, but it needs to be confirmed.We illustrated a possible pathway that involved in *CRTC1* KD induced cell-death signals, followed by aberrant IEG expression (Figure S10). In addition to the Figure S10 legend, *de novo* SIK1 mutations cause SIK syndrome, impacting HDAC5, synaptic activity response genes in humans and mice^25–27^. *SIK1* variation disrupts *MEF2*, *Nur77* (NR4A1) and *Neureglin 1* expressions^25^. *Creb* and *Crem* disruption in mice leads to apoptosis in postmitotic neurons and postnatal forebrain, causing hippocampal, striatal neurodegeneration^28^. Correlated roles of *Sirt1*, *Crtc1*, *CREB* in the neuroprotective pathway of mutant mouse huntingtin are reported^29^.

The marmoset epilepsy model based on the current studies may have the advantage of analyzing epilepsy by ECoG and dMRI in real time from the onset of the event. Epilepsy is typically manifested by recurrent seizures and can only be diagnosed at unavoidable intervals. Unlike previous models which were induced by trauma, infection, ischemia-induced membrane potential changes or brain damage^7^, channel genes^30^, mice deficient in all three PAR bZip proteins^31^, and a family of marmosets with generalized seizures which was measured by low frequency oscillation (LOF) induced by handling operation^32^, the *shCRTC1* model uses single gene local KD. This allowed us to observe the chronological progression of epilepsy, as *shCRTC1* KD marmosets revealed the early mechanisms of epilepsy. Thus, this approach may lead to the development of new antiepileptic drugs targeting early-onset epilepsy. It could also be applied to early diagnosis, anticonvulsant prescription and treatment of focal and structural epilepsy.

## Limitation of the study

A potential limitation of the current study is the off-target effect of *shCRTC1*#1 on epilepsy. However, despite the differences between the five shCRTC1s, this possibility is unlikely for several reasons, although we cannot completely exclude it. First, these *shCRTC1*s are highly selective for the *CRTC1* gene in the marmoset genome (see STAR METHODS shRNA preparation and plasmid construction, and Figure S9b). Second, although *shCRTC1*#2 tends to induce more cell death than #1, they also share a similar function of inducing IEG, at least to some extent. Furthermore, Monkey 9, injected with *shCRTC1*#1 at one site and *shCRTC1*#2 and #3 at one and two sites, respectively, showed a wider IEG expression similar to monkey 1 (Supplemental Table 1a). It remains for future studies to know how precisely each different *shCRTC1* works and interacts with each other.

Another limitation of the present study, related to the above potential limitation, is that the causal relationship between cell death and IEG induction remains unclear. The relationship may be interdependent. That is, the delicate balance between the cell death pathway and increased neuronal activity may explain why *shCRTC1*#1 injections were directed toward the epileptic response, whereas *shCRTC1*#2 injections were directed toward the cell death pathway with the slower IEG induction than *shCRTC1*#1. This delicate balance may also explain the difference in IEG induction between Monkeys 4 and 9 without TTX and with monocular TTX injections in the retina when shCRTC1#1 was injected at a single site in the V1 hemisphere. In any case, the cellular damage caused by the local injection of *shCRTC1*s in V1 converged on remission at the individual animal level, but the pathways were quite different for the two *shCRTC1* types. One led to a cortical-wide epileptic response, while the other led to localized cellular damage. Further investigation of how the different *shCRTC1*s act at the molecular and cellular level may provide clues to understanding the balance and critical points of these two pathways.

## Acknowledgements

We thank RIKEN RRD and N. Hasegawa for their support. We also thank Dr. Hirofumi Nakatomi for suggestions on various aspects of epilepsy studies. This work was supported by the Program for Scientific Research on Innovative Areas (grant number, 22123009) from MEXT, Japan, and by the Program for Brain Mapping by Integrated Neuro technologies for Disease Studies (Brain/MINDS: JP15dm0207001 to T.Y. and M.K., JP19dm0207069 to M.K., JP19dm0207088 to K.N., JP15dm0207001 to H.O.) from AMED. JSPS KAKENHI (JP22H05154 and 22H05163 to K.N., JP22H03630 to J.H.)

## Contributions

Y.N., M.K., K.N., and T.Y. designed the study and drafted the manuscript. Y.N., M.O., and A.W. performed the virus injections and histological studies. H.T. provided the anti-CRTC1 antibody. H.T., M.O., and A.W. participated in the discussion of CRTC1 function. M.K. established the ECoG system. M.K. and Y. N. performed the ECoG experiments. M. K. and K.N. analyzed the ECoG and dMRI data. J.H. collected the dMRI data and analyzed the dMRI data. H.M. prepared the AAV vectors. S.I., H.O., and T.Y. supported and supervised the project.

## Competing interests

All authors declare no competing interests. RIKEN holds the patent (2022-15863) for this study in Japan.

## METHODS

### Experimental procedures and marmosets used

The experimental procedures, including tissue preparation, *in situ* hybridization (ISH), and immunohistochemistry (IHC), were essentially performed as described or cited in our previous work unless specifically stated otherwise below. The earlier experiments were conducted in accordance with the guidelines of the National Institutes of Health and the Ministry of Education, Culture, Sports, Science and Technology (MEXT) of Japan and were approved by the Animal Care and Use Committee of the National Institutes of Natural Sciences. All other experimental procedures were approved by the Experimental Animal Committee of RIKEN.

### shRNA preparation and plasmid construction

All the shRNA sequences were designed using siRNA Sequence Selector (Clontech). AAV vectors for shRNA expression (see Figure 1a) were based on pAAV-U6-shRNA-2A-hSYNI-hrGFP^33^. A shRNA cassette was inserted into the plasmid digested with BamHI and HindIII located directly below the U6 promoter. To detect the infected cell, hSYNI-hrGFP was replaced Sirius^34^ fused human histone H2B (Gene ID: 8970, GenBank ID: X00088) with 3xFlag tag driven by the short version of the mouse CamKIIa promoter. Sirius/pcDNA3 was a gift from Dr. Takeharu Nagai (Addgene http://n2t.net/addgene: plasmid #51957; RRID: Addgene_51957).

### Manipulation procedure for each marmoset used

A total of 10 adult common marmosets (*Callithrix jacchus*, 14–69 months, both sexes) were used in this study (Supplemental Table 1a). Unless otherwise stated, the marmosets were normally reared (NR) under 12hr light and 12hr dark conditions. Perfusion was performed after the animals had been under the light cycle for more than 6hr. Eight marmosets except for Monkeys 1 and 6 did not receive TTX injections and were therefore visually intact. TTX injection and visual manipulation were performed as described previously^11^ with minor modifications. Monkey 1 was kept in the dark for 24–41hr after TTX injection into one eye, followed by LS for 24 min before sacrifice. The experimental schedule and shRNA injection sites of Monkey 1 are shown in Figures 1b and S3a. Monkeys 2 and 3 were used for ECoG recording after injection of *shCRTC1*#1 at three sites in the right hemisphere. Monkey 4 was normally reared and the injection sites are shown in Figure S8a. Monkeys 5 and 6 were used for light induction (LS) experiments as follows: Monkey 5 was kept in the dark for 24–41hr, followed by LS for 24 min before sacrifice and fixation. Monkey 6 was kept in the dark for 24–41hr after TTX injection into one eye, followed by LS for 24 min before sacrifice (Supplemental Table 1a). Monkey 7 was normally reared and injected with each of *shscr*, *shCRTC*1#2 and #3 in both hemispheres. The injection sites for Monkeys 8, and 10 are shown in and Figure S9h and Figure 7a, respectively. Monkey 9 was normally reared and the injection numbers of each *shCRTC1* and *shscr* in both hemispheres are shown in Supplemental Table 1a.

### AAV injection

Adeno-Associated Virus serotype 1 was used in all the experiments. They were produced in HEK293T cells^33, 35, 36^, the line used in the previous publication, using a helper virus-free system, and purified twice with CsCl2 density gradients and titrated by Q-PCR as described previously^37^. Surgery for *shRNA* injection was performed as previously described with some modifications^38^. Stereotaxic positions were aligned by using the interaural plane and the anterior border of the cortex. The injection position was determined by the distance from the posterior end of the exposed head bone for the AP direction, and by the distance from the midline of the skull for the ML direction. All injections were performed in V1 on the cortical surface, and adjacent injection sites were injected 2-3 mm apart in the DM and ML directions, respectively. Pressure injection was performed using a 30 μm outer diameter glass micropipette connected to a nanoliter 2000 injector with a Micro4 controller (World Precision Instruments). To inject AAV into the marmoset V1 cortex, the injector was tilted 30 degrees to the tangent perpendicular line. For exposed cortical areas, we injected 0.1 μl each of vector [5×10^9^ viral genome (vg)/μl] at two depths (0.3-0.5 and 0.6-1.1 mm from the surface). Since the injector was tilted, the depth was varied according to the position of the injection site, with the aim of delivering AAV to all cortical layers. The injection needle was placed at a shallow level in the cortex, left in place for 2 min, and then an injection of 0.1 μl/2.5 min was delivered. The needle was then advanced further, left in place for 2 min, and a second injection of 0.1 μl/2.5 min was given. After a further waiting 2 min, the needle was removed. Note that in Monkey 5, when *shCRTC1*#1 was injected into the left hemisphere, the glass needle was not aligned with the surface of the brain and it was not injected deeply.

### ISH

The ISH method was performed on free-floating sections as previously described^39^. Briefly, brain sections (25 μm) were pretreated with 0.1 M PB, 0.75% glycine/0.1 M PB, and 0.3% Triton X-100/0.1 M PB followed by protease K treatment (5 mg, 37°C, 30 min). After acetylation at RT for 10 min, prehybridization and hybridization were performed at 60-70°C for 1hr and overnight, respectively. Sections were washed with 2x standard saline citrate (SSC)/50% formamide/0.1% N-lauroylsarcosine (NLS), treated with RNase A (20 μg/ml), washed again with 2xSSC/50% formamide/0.1% NLS, and immersed in blocking solution (60 min, RT). For single-color ISH, signals were detected by antibody reaction with anti-digoxigenin (DIG)-AP, Fab fragment (Merck) followed by the nitro blue tetrazolium chloride (NBT)/5-bromo-4-chloro-3-indolyl-phosphate (BCIP) system. For two-color ISH, the DIG probe was detected using an HNPP fluorescent detection kit (11758888001, Merck). The FITC probe was detected as follows. The antibody reaction was performed with Peroxidase-IgG Fraction Monoclonal Mouse Anti-FITC (#200-032-037, Jackson ImmunoResearch Laboratory). The signal was then amplified using a TSA PLUS system (NEL747A001KT, Akoya) and detected with Alexa Fluor 488-conjugated anti-DNP antibody (Molecular Probes). The ISH probes are listed in Supplemental Table 1b. The *cFOS, ZIF268*, *AR*C, *HTR1B* and *HTR2A* probes and the contents and procedures for each reaction used were described in our previous publication^39^.

### Immunohistochemistry (IHC)

IHC was performed using a free-floating section method as previously described. Briefly, brain sections were washed with Tris-buffered saline (TBS). The sections were blocked at RT in blocking buffer containing 5% bovine serum albumin, 0.1% Triton X 100, and 4% normal goat serum in TBS, followed by an antibody reaction with the primary antibody at 4°C for 16-72hr. After the samples were washed with TNT buffer (0.1 M Tris-HCl, pH 7.5, 0.15 M NaCl, 0.1% Tween 20), the secondary antibody reaction was performed at RT for 2hr. For multiple fluorescent labeling, the sections were washed with TNT buffer, mounted on glass slides, and embedded in gelatin. For single-color staining, the sections were labeled with avidin-biotin complex using a Vectastain ABC Elite kit (Vector Laboratories, Burlingame, CA) at RT for 1hr, stained with DAB, and embedded in glass slides. Primary antibodies were TORC1/CRTC1 (C71D11) rabbit mAb (2587, Cell Signaling Technology, Inc.), anti-phospho-CREB (Ser133) antibody, clone 10E9 (05-667, Merck), anti-GFAP antibody (ab7260, Abcam), anti-Iba1 rabbit antibody (019-19741, FUJIFILM Wako), anti-NeuN antibody (MAB377, Merck), and anti-c-Fos rabbit polyclonal IgG antibody (sc-52, Santa Cruz Biotechnology). Because the fluorescence signal of the Sirius protein in brain sections was weak, the signal was enhanced by antibody staining with anti-GFP chicken antibody (ab13970, Abcam), unless otherwise noted. Secondary antibodies were biotinylated (Jackson ImmunoResearch Laboratories) and fluorophore-conjugated antibodies [CyTM2, CyTM3, CyTM5 (Jackson ImmunoResearch Laboratories), and Alexa Fluor 488 (Molecular Probes)].

### Cell death assay (TUNEL staining)

TUNEL staining was performed using the *In Situ* Apoptosis Detection Kit (TAKARA BIO INC., Japan) with minor modifications. Briefly, free-floating sections were postfixed in 4% paraformaldehyde (PFA) in PB for 2hr at RT or overnight at 4°C. Sections were sequentially treated with 0.75% glycine, 0.3% Triton X-100, 5 mg/ml protease K (incubated at 37°C for 30 min) and 1% H_2_O_2_ in phosphate-buffered saline (PBS) (at RT for 30 min). The following procedures were performed according to the manufacturer’s protocol. After staining with DAB, the sections were mounted on gelatin-coated slides, and Nissl staining was performed as a counterstain, if required.

### Histological data Processing

The histological images were obtained using a digital color camera DP70 (Olympus, Tokyo, Japan) attached to a BX-51 microscope (Olympus,Tokyo, Japan), and an all-in-one microscope (Keyence BZ-X710). After image acquisition of brain sections using microscopes, image contrast was optimized, and false colors were applied using Photoshop CS5 or Photoshop 2022 software (Adobe Systems, San Jose, CA) according to AMED image processing guidelines (https://www.amed.go.jp/content/000078447.pdf). Although the sections for ISH were shrunk, the scale bars in the figures indicate the size of the mounted sections, which were not adjusted for shrinkage.

### Large Language Models (LLMs)

We used ChatGPT (https://chat.openai.com) to examine some text sentences, writing only on the draft that were written by the authors. We did not use it for the current data analysis.

### ECoG recordings

Whole-cortex 64-channel and 96-channel ECoG arrays (Cir-Tech Inc., Japan) were implanted epidurally in the right hemisphere of two monkeys (Monkeys 2 and 3). Ten electrodes implanted in Monkey 2 were cut during implantation. The implantation procedure has been described in detail previously^12^. At the time of implantation (Monkey 2) or 1 week prior to implantation (Monkey 3), the *CRTC1* KD AAV vector was injected into three sites in V1 of the right hemisphere of the monkeys (Figure 5a for Monkey 2). Longitudinal ECoG recordings and behavioral monitoring were conducted for 2 and 5 months after the AAV injections in Monkeys 2 and 3, respectively. During daily 1-2hr monitoring sessions, the monkeys were seated in a primate chair in a dimly lit room. For the first 20 min of the recordings, we presented sound stimuli used in a previous study^14^ to examine auditory evoked responses. ECoG signals were recorded at a sampling rate of 1 kHz per channel by using a Cerebus system (Blackrock Microsystems, USA).

### Anatomical localization of electrode contacts

The positions of each electrode contact were identified based on postoperative computed tomography and preacquired T2-weighted MRI. The localization procedure has been described in detail previously (Brain/MINDS data portal; DataID: 4958). Based on the putative cortical areas, we divided the electrodes into seven groups, consisting of V1 and visual (Vis), auditory (Au), temporal (TC), sensorimotor (SM), posterior parietal (PC), and prefrontal areas (FC). Vis was defined as visual areas excluding V1, the site of KD injections. In all monkeys, the electrode array covered the frontal, parietal, occipital, and temporal cortices (Figure 2b).

### dMRI data acquisition

*Ex vivo* brains were obtained after transcardial perfusion with 4% paraformaldehyde in 0.1 M phosphate buffer (pH 7.4), the marmoset brains were removed, post-fixed at 4° C for 2-3 days, and transferred to 50 mM phosphate buffer (pH 7.4), and scanned 2-3 days after fixation. All marmoset brains had an *ex vivo* MRI scan (T2w and DWI) before further processing. MRI was performed using a 9.4-T BioSpec 94/30 unit (Bruker Optik GmbH) and a transmit and receive solenoid type coil with a 28-mm inner diameter. To acquire *ex vivo* brain data, the brain was wrapped in a sponge and soaked in a fluorine solution, which showed no signal on MRI, in a plastic container. Vacuum degassing was performed to reduce air bubble-derived artifacts. dMRI data sets were acquired using a spin‒echo sequence based on Stejskal-Tanner diffusion^40^. Scanning parameters were as follows: repetition time (TR), 4000 ms; echo time (TE), 28.4 ms; flip angle, 90°; field of view (FOV), 38×38 mm; acquisition data matrix, 190×190; reconstructed image resolution, 0.2 mm (with zero-filling interpolation); slice thickness, 0.2 mm; b-value, 3000 s/mm^2^; motion-probing gradient (MPG) orientations, 128 axes; and number of averages (NA), 2.

### Data analysis

After preprocessing, quantitative analysis was performed. For quantitative analysis, the ECoG signal was clipped to the first 15-min recording of each experimental day. The signals were then rereferenced using a common mean reference (CMR) montage (Figures S4a and S5a). In this process, we rereferenced the signal from each channel to the common mean signal across all channels, from which the channel(s) with a standard deviation greater than 250 mV were excluded. The signals were used to calculate functional connectivity. To extract HFA, we applied a bandpass filter (80 to 200 Hz) to the signal and then calculated the envelope using the Hilbert transform (Figures S4b and S5b).

### Detection of cwHFO

We defined an HFO as an HFA greater than the mean plus two times the standard deviation of the HFA across all channels. We then counted the number of electrodes showing HFOs at each time point (Figures S4c and S5c), and an event was marked as a cwHFO if the proportion of electrodes exceeded 50% (=0.5). Marked events with an interevent interval of less than 1 s were merged into one cwHFO. During some putative cwHFOs, Monkey 5 happened to bite the chair, and it was difficult to separate the biting-related noise from neuronal activity. Therefore, we rejected the events from the putative cwHFOs by using a supervised learning method. We recorded movies from marmosets on seven days (22, 35, 42, 51, 55, 57, and 62 days after KD). Using these movies, we manually distinguished biting events from the putative cwHFOs. We labeled 39 biting events from out of 296 cwHFOs. We then trained a linear support vector machine (SVM) to classify these cwHFOs into biting events and others. We focused on the time-averaged HFA of the 33 electrodes that were stably observed on all experimental days. The SVM was applied to the time-averaged signal for time windows ranging from 0 to 400 ms relative to the onset of the cwHFOs. Note that this time window size was optimized using 10-fold cross validation over the seven days. We found that 94% of the cwHFOs occurred on days when we did not record movies and removed the events labeled as biting events from the following analysis. To investigate which cortical region was the origin of the cwHFO, we calculated the mean HFA for the seven cortical groups in a time window ranging from -250 to 0 ms for each cwHFO (V1, Vis, Au, TC, SM, PC, and FC).

### Functional connectivity

We obtained a functional connectivity matrix for each recording by using the correlation coefficient from within 15 min of ECoG signals from each electrode. We then averaged all the matrices and compared them with each daily connectivity matrix to evaluate the daily deviation from the average. To examine the day-to-day stability of the functional connectivity, we calculated (1) correlations among the connectivity matrices on experimental days and (2) the mean functional connectivity within and between regions. Finally, to visualize changes in the connectivity within the KD site (V1) for both monkeys, we calculated the p value of the correlation coefficients within the V1 signals among the experimental days using Fisher’s z transformation. The transformed correlation coefficients follow a normal distribution under regular conditions^41^.

We spatially interpolated the HFA of each electrode to the surface of the standard marmoset brain, BMA 2019 Ex vivo^42^. We construct the left and right surfaces of the standard brain using iso2Mesh^43^, where the number of hemisphere vertices is 1672 and the average distance between the vertices is 1 mm. The interpolated HFA of each vertex at a specific timepoint was calculated using a Gaussian process regression^44^ with a 3-dimensional isotropic Gaussian kernel of sigma = 4 mm.

### dMRI analysis

We obtained structural connectivity maps based on 36 individual *ex vivo* dMRI scans by using a global tractography method^44^. We used the MRI of a standard brain from the Brain/MINDS portal (https://doi.org/10.24475/bma.4520), whose left and right mid-surface was defined on 167082 vertices. We randomly sampled the 12428 vertices from the original surface and defined corresponding supervoxels on the standard brain. The acquired diffusion-weighted MRI data were analyzed to perform DTT using Mrtrix3 and Diffusion Toolkit software (Massachusetts General Hospital, Boston, MA, USA). Tractography was performed based on fiber assignments using a continuous tracking algorithm^45^. We obtained a connectivity matrix among the supervoxels (7.2 mm^3^) to count the number of streamlines through two supervoxels. We focused on the asymmetricity of the convexity pattern between the left and right cortices. By setting two symmetric points on the left and right cortices, we calculated the correlation between the connectivity profiles of each cortex at the two points. A low correlation indicates an asymmetric connectivity pattern at these points (the red region of Figure. 4a). To evaluate the asymmetric feature of the epilepsy-model marmosets, we calculated the z score of the asymmetric connectivity pattern of normal marmosets averaged over brain regions, which we refer to as the asymmetric feature (AF) in Fig. 4b. Fibers passing through the V1 and frontal areas or the V1 and temporal areas were labeled V1-F and V1-T, respectively. Fibers were counted for each right and left hemisphere. We calculated the asymmetry rates on dMRI as R/(R+L), where R and L correspond to the fiber counts for the right and left hemispheres, respectively.

## Notes

### Competing Interest Statement

The authors have declared no competing interest.

### Summary of Updates

The main point of this revision is that the order of presentation of the figures is largely changed as follows, 1) We first described the results with three injection sites of shCRTC1#1 in V1, which caused a high level of IEG including cFOS expressions and epileptic responses. 2) We then described the effect with fewer injection sites and/or less potent shCRTC1s than shCRTC1#1, although the short hairpin (sh) sequences perfectly match the marmoset CRTC1 RNA. 3) Under the fewer injection sites and/or less potent shCRTC1s, IEG expression is weak, but we observed significant cell lesions and expression of cell death related markers such as IBA1 and GFAP. These results suggest that the mutual interaction between cell death signals and abnormal enhancement of neuronal activity, which eventually cause epileptic response in the brain.

## References

1. Watanabe, S., Kurotani, T., Oga, T., Noguchi, J., Isoda, R., Nakagami, A., Sakai, K., Nakagaki, K., Sumida, K., Hoshino, K., et al. (2021). Functional and molecular characterization of a non-human primate model of autism spectrum disorder shows similarity with the human disease. Nat Commun 12, 5388. 10.1038/s41467-021-25487-6.

2. Schmidt, D., and Sillanpaa, M. (2012). Evidence-based review on the natural history of the epilepsies. Curr Opin Neurol 25, 159–163. 10.1097/WCO.0b013e3283507e73.

3. Bialer, M., and White, H.S. (2010). Key factors in the discovery and development of new antiepileptic drugs. Nat Rev Drug Discov 9, 68–82. 10.1038/nrd2997.

4. White, H.S. (2003). Preclinical development of antiepileptic drugs: past, present, and future directions. Epilepsia 44 Suppl 7, 2–8. 10.1046/j.1528-1157.44.s7.10.x.

5. Depaulis, A., David, O., and Charpier, S. (2016). The genetic absence epilepsy rat from Strasbourg as a model to decipher the neuronal and network mechanisms of generalized idiopathic epilepsies. J Neurosci Methods 260, 159–174. 10.1016/j.jneumeth.2015.05.022.

6. van Luijtelaar, G., and Zobeiri, M. (2014). Progress and outlooks in a genetic absence epilepsy model (WAG/Rij). Curr Med Chem 21, 704–721. 10.2174/0929867320666131119152913.

7. Pitkanen, A., Kharatishvili, I., Karhunen, H., Lukasiuk, K., Immonen, R., Nairismagi, J., Grohn, O., and Nissinen, J. (2007). Epileptogenesis in experimental models. Epilepsia 48 *Suppl 2*, 13–20. 10.1111/j.1528-1167.2007.01063.x.

8. Loscher, W., Klitgaard, H., Twyman, R.E., and Schmidt, D. (2013). New avenues for anti-epileptic drug discovery and development. Nat Rev Drug Discov 12, 757–776. 10.1038/nrd4126.

9. Berg, A.T., Berkovic, S.F., Brodie, M.J., Buchhalter, J., Cross, J.H., van Emde Boas, W., Engel, J., French, J., Glauser, T.A., Mathern, G.W., et al. (2010). Revised terminology and concepts for organization of seizures and epilepsies: report of the ILAE Commission on Classification and Terminology, 2005-2009. Epilepsia 51, 676–685. 10.1111/j.1528-1167.2010.02522.x.

10. Scheffer, I.E., Berkovic, S., Capovilla, G., Connolly, M.B., French, J., Guilhoto, L., Hirsch, E., Jain, S., Mathern, G.W., Moshe, S.L., et al. (2017). ILAE classification of the epilepsies: Position paper of the ILAE Commission for Classification and Terminology. Epilepsia 58, 512–521. 10.1111/epi.13709.

11. Nakagami, Y., Watakabe, A., and Yamamori, T. (2013). Monocular inhibition reveals temporal and spatial changes in gene expression in the primary visual cortex of marmoset. Frontiers in Neural Circuits 7. 10.3389/fncir.2013.00043.

12. Komatsu, M., Takaura, K., and Fujii, N. (2015). Mismatch negativity in common marmosets: Whole-cortical recordings with multi-channel electrocorticograms. Sci Rep 5, 15006. 10.1038/srep15006.

13. Kaneko, T., Komatsu, M., Yamamori, T., Ichinohe, N., and Okano, H. (2022). Cortical neural dynamics unveil the rhythm of natural visual behavior in marmosets. Commun Biol 5, 108. 10.1038/s42003-022-03052-1.

14. Jiang, Y., Komatsu, M., Chen, Y., Xie, R., Zhang, K., Xia, Y., Gui, P., Liang, Z., and Wang, L. (2022). Constructing the hierarchy of predictive auditory sequences in the marmoset brain. Elife 11. 10.7554/eLife.74653.

15. Watakabe, A., Ichinohe, N., Ohsawa, S., Hashikawa, T., Komatsu, Y., Rockland, K.S., and Yamamori, T. (2007). Comparative analysis of layer-specific genes in mammalian neocortex. Cerebral Cortex 17, 1918–1933. 10.1093/cercor/bhl102.

16. Burnos, S., Hilfiker, P., Surucu, O., Scholkmann, F., Krayenbuhl, N., Grunwald, T., and Sarnthein, J. (2014). Human intracranial high frequency oscillations (HFOs) detected by automatic time-frequency analysis. PLoS One 9, e94381. 10.1371/journal.pone.0094381.

17. Komatsu, M., Kaneko, T., Okano, H., and Ichinohe, N. (2019). Chronic Implantation of Whole-cortical Electrocorticographic Array in the Common Marmoset. J Vis Exp. 10.3791/58980.

18. Jacobs, J., Staba, R., Asano, E., Otsubo, H., Wu, J.Y., Zijlmans, M., Mohamed, I., Kahane, P., Dubeau, F., Navarro, V., and Gotman, J. (2012). High-frequency oscillations (HFOs) in clinical epilepsy. Prog Neurobiol 98, 302–315. 10.1016/j.pneurobio.2012.03.001.

19. Jirsch, J.D., Urrestarazu, E., LeVan, P., Olivier, A., Dubeau, F., and Gotman, J. (2006). High-frequency oscillations during human focal seizures. Brain 129, 1593–1608. 10.1093/brain/awl085.

20. Devinsky, O., Vezzani, A., Najjar, S., De Lanerolle, N.C., and Rogawski, M.A. (2013). Glia and epilepsy: excitability and inflammation. Trends Neurosci 36, 1784. 10.1016/j.tins.2012.11.008.

21. Loewen, J.L., Barker-Haliski, M.L., Dahle, E.J., White, H.S., and Wilcox, K.S. (2016). Neuronal Injury, Gliosis, and Glial Proliferation in Two Models of Temporal Lobe Epilepsy. J Neuropathol Exp Neurol 75, 366–378. 10.1093/jnen/nlw008.

22. Nonaka, M., Kim, R., Fukushima, H., Sasaki, K., Suzuki, K., Okamura, M., Ishii, Y., Kawashima, T., Kamijo, S., Takemoto-Kimura, S., et al. (2014). Region-specific activation of CRTC1-CREB signaling mediates long-term fear memory. Neuron 84, 92–106. 10.1016/j.neuron.2014.08.049.

23. Xu, S., Sun, Q., Fan, J., Jiang, Y., Yang, W., Cui, Y., Yu, Z., Jiang, H., and Li, B. (2019). Role of Astrocytes in Post-traumatic Epilepsy. Front Neurol 10, 1149. 10.3389/fneur.2019.01149.

24. Sano, F., Shigetomi, E., Shinozaki, Y., Tsuzukiyama, H., Saito, K., Mikoshiba, K., Horiuchi, H., Cheung, D.L., Nabekura, J., Sugita, K., et al. (2021). Reactive astrocyte-driven epileptogenesis is induced by microglia initially activated following status epilepticus. JCI Insight 6. 10.1172/jci.insight.135391.

25. Proschel, C., Hansen, J.N., Ali, A., Tuttle, E., Lacagnina, M., Buscaglia, G., Halterman, M.W., and Paciorkowski, A.R. (2017). Epilepsy-causing sequence variations in SIK1 disrupt synaptic activity response gene expression and affect neuronal morphology. Eur J Hum Genet 25, 216–221. 10.1038/ejhg.2016.145.

26. Hansen, J., Snow, C., Tuttle, E., Ghoneim, D.H., Yang, C.S., Spencer, A., Gunter, S.A., Smyser, C.D., Gurnett, C.A., Shinawi, M., et al. (2015). De novo mutations in SIK1 cause a spectrum of developmental epilepsies. Am J Hum Genet 96, 682–690. 10.1016/j.ajhg.2015.02.013.

27. Pang, B., Mori, T., Badawi, M., Zhou, M., Guo, Q., Suzuki-Kouyama, E., Yanagawa, T., Shirai, Y., and Tabuchi, K. (2022). An Epilepsy-Associated Mutation of Salt-Inducible Kinase 1 Increases the Susceptibility to Epileptic Seizures and Interferes with Adrenocorticotropic Hormone Therapy for Infantile Spasms in Mice. Int J Mol Sci 23. 10.3390/ijms23147927.

28. Mantamadiotis, T., Lemberger, T., Bleckmann, S.C., Kern, H., Kretz, O., Martin Villalba, A., Tronche, F., Kellendonk, C., Gau, D., Kapfhammer, J., et al. (2002). Disruption of CREB function in brain leads to neurodegeneration. Nat Genet 31, 47–54. 10.1038/ng882.

29. Jeong, H., Cohen, D.E., Cui, L., Supinski, A., Savas, J.N., Mazzulli, J.R., Yates, J.R., 3rd, Bordone, L., Guarente, L., and Krainc, D. (2011). Sirt1 mediates neuroprotection from mutant huntingtin by activation of the TORC1 and CREB transcriptional pathway. Nat Med 18, 159–165. 10.1038/nm.2559.

30. Oyrer, J., Maljevic, S., Scheffer, I.E., Berkovic, S.F., Petrou, S., and Reid, C.A. (2018). Ion Channels in Genetic Epilepsy: From Genes and Mechanisms to Disease-Targeted Therapies. Pharmacol Rev 70, 142–173. 10.1124/pr.117.014456.

31. Gachon, F., Fonjallaz, P., Damiola, F., Gos, P., Kodama, T., Zakany, J., Duboule, D., Petit, B., Tafti, M., and Schibler, U. (2004). The loss of circadian PAR bZip transcription factors results in epilepsy. Genes Dev 18, 1397–1412. 10.1101/gad.301404.

32. Yang, X., Chen, Z., Wang, Z., He, G., Li, Z., Shi, Y., Gong, N., Zhao, B., Kuang, Y., Takahashi, E., and Li, W. (2022). A natural marmoset model of genetic generalized epilepsy. Mol Brain 15, 16. 10.1186/s13041-022-00901-2.

33. Takaji, M., Takemoto, A., Yokoyama, C., Watakabe, A., Mizukami, H., Ozawa, K., Onoe, H., Nakamura, K., and Yamamori, T. (2016). Distinct roles for primate caudate dopamine D1 and D2 receptors in visual discrimination learning revealed using shRNA knockdown. Sci Rep 6, 35809. 10.1038/srep35809.

34. Tomosugi, W., Matsuda, T., Tani, T., Nemoto, T., Kotera, I., Saito, K., Horikawa, K., and Nagai, T. (2009). An ultramarine fluorescent protein with increased photostability and pH insensitivity. Nat Methods 6, 351–353. 10.1038/nmeth.1317.

35. Watakabe, A., Ohsawa, S., Hashikawa, T., and Yamamori, T. (2006). Binding and complementary expression patterns of semaphorin 3E and plexin D1 in the mature neocortices of mice and monkeys. J Comp Neurol 499, 258–273. 10.1002/cne.21106.

36. Hata, K., Mizukami, H., Sadakane, O., Watakabe, A., Ohtsuka, M., Takaji, M., Kinoshita, M., Isa, T., Ozawa, K., and Yamamori, T. (2013). DNA methylation and methyl-binding proteins control differential gene expression in distinct cortical areas of macaque monkey. J Neurosci 33, 19704–19714. 10.1523/JNEUROSCI.2355-13.2013.

37. Yagi, H., Ogura, T., Mizukami, H., Urabe, M., Hamada, H., Yoshikawa, H., Ozawa, K., and Kume, A. (2011). Complete restoration of phenylalanine oxidation in phenylketonuria mouse by a self-complementary adeno-associated virus vector. J Gene Med 13, 114–122. 10.1002/jgm.1543.

38. Watakabe, A., Skibbe, H., Nakae, K., Abe, H., Ichinohe, N., Rachmadi, M.F., Wang, J., Takaji, M., Mizukami, H., Woodward, A., et al. (2023). Local and long-distance organization of prefrontal cortex circuits in the marmoset brain. Neuron 111, 2258–2273 e2210. 10.1016/j.neuron.2023.04.028.

39. Watakabe, A., Ichinohe, N., Ohsawa, S., Hashikawa, T., Komatsu, Y., Rockland, K.S., and Yamamori, T. (2007). Comparative analysis of layer-specific genes in Mammalian neocortex. Cereb Cortex 17, 1918–1933. 10.1093/cercor/bhl102.

40. Stejskal, E.O., Tanner, J.E. (1965). Spin diffusion measurements: Spin echoes in the presence of a time-dependent field gradient. J. Chem. Phys. 42, 288–292.

41. Fisher, R.A. (1915). Frequency distribution of the values of the correlation coefficient in samples from an indefinitely large population. Biometrika 10.*4*, 507–521.

42. Woodward, A., et al. (2020). The NanoZoomer artificial intelligence connectomics pipeline for tracer injection studies of the marmoset brain. Brain Struct Funct 225, 1225–1243.

43. Tran, A.P., Yan, S., and Fang, Q. (2020). Improving model-based functional near-infrared spectroscopy analysis using mesh-based anatomical and light-transport models. Neurophotonics 7, 015008. 10.1117/1.NPh.7.1.015008.

44. Schulz, E., Speekenbrink, M., & Krause, A. (2018). A tutorial on Gaussian process regression: Modelling, exploring, and exploiting functions. Journal of Mathematical Psychology, 85, 1–16. 10.1016/j.jmp.2018.03.001

45. Mori, S., and van Zijl, P.C. (2002). Fiber tracking: principles and strategies - a technical review. NMR Biomed 15, 468–480. 10.1002/nbm.781.

