## Supplemental legends for "Epileptic responses in marmosets induced by knockdown of CREB Regulated Transcription Coactivator 1 (CRTC1)"

**Figure S1. *shCRTC1* construction and its effect on Nurr77 (continued from Fig. 1). a,** Construction of an expression vector for the marmoset *CRTC1* gene (left). Western blot (WB) analysis (right). The specificity of *shCRTC1*#1 and #2 was examined by cotransfecting each of the *shCRTC1*s together with the marmoset *CRTC1* gene into HEK293T cells. Each WB column shows cotransfection with no shRNA, *shscr*, *shCRTC1*#1, and s*hCRTC1*#2 (from left to right). Each row shows detection with anti-CRTC1, anti-HA and anti-β-actin (from top to bottom). **b,** Top two rows: IHC analysis at the *shCRTC*#1 and *shscr* injection sites in marmoset V1 detected with anti-Sirius (GFP) antibody (top) and CRTC1 antibody (bottom). Adjusted sections were analyzed. Scale bar (1 mm). The bottom three right rows are magnified views of the white square regions (a and b in the small white boxes) in the CRTC1 section, showing Sirius fluorescence, anti-CRTC1 IHC signals, and merged images (top, middle, and bottom rows, respectively) at the *shCRTC*1#1 and *shscr* injection sites (right and left columns, respectively). **c,** Images from ISH analysis showing the *NUR77* signal alone and merged with the Sirius signal (from left to right) at the *shCRTC1*#1 injection site. Layers 1, 2, 4C, 5, 6 and white matter are shown (L1, L2, L4C, L5, L6, and WM, respectively). **d,** ISH analysis of *NURR1* with *Sirius*. Top: *NURR1* alone and merged with *Sirius* (from left to right) at the *shCRTC1*#1 noninjection and injection sites indicated by white boxes 1 and 2, respectively. Layers indicated by L1, L2, L6a and L6b. Bottom: Enlarged views of the above white boxes (box 1, left, and box 2, right) showing layers 6a (L6a) and 6b (L6b). Scale bar (100 μm).

**Figure S2. IEG expression around *shCRTC1* and shscr injection sites.** Left column: Four IEG expression patterns were examined around the *shCRTC1* and *shscr* injection sites in Monkey 1. Middle column: Sirius ISH. Right column: Merged images of the IEG and Sirius signals. Rows from top to bottom: *cFOS*, *ARC*, *ZIF268*, and *NURR1* expression by ISH. Arrowheads indicate the border of each IEG expression. Scale bar: 500 μm.

**Figure S3: Neuronal, glial and cell death markers around the shRNA injection site in Monkey 1. a,** Top left: Illustration of the injection sites. The cross-section, as shown by the dotted line, which includes the *shCRTC1*#1 and *shscr* injection sites, was analyzed histologically. Bottom left: Nissl staining around an injection site. Histological analysis of Nissl, glial, neuronal and cell death markers. Middle and right panels: Double IHCs of glial markers and Sirius. IBA1 (middle column) and GFAP (right column) signals are shown in magenta. Sirius (green) was used for injection site detection. Adjusted sections were analyzed. Nissl staining was performed on the same section as IBA1 immunostaining. Scale bar 500 μm. **b**, Left and middle: NeuN and Sirius staining are shown in magenta and green, respectively. Right: merged images. Scale bar 200 μm. **c,** TUNEL staining at the injection site. Top left panel is an image of the boxed region of the right panel (enlarged TUNEL-stained region). Bottom left: Sirius. Scale bar 500 μm. Right: Low-magnification view of the *shCRTC1*-injected region. **d,** Top and middle: Sirius expression and phosphorylated CREB signals, were detected at the *shCRTC1* injection site, respectively. Bottom: CRTC1 expression (IHC).

**Figure S4. ECoG signals from Monkey 2. a.** Examples of raw ECoG signals for 2 minutes. The x-axis is time, and the y-axis is the electrode alignment. The amplitude (μv) is also shown in the time scale (s) in the lower right corner of each panel. **b.** Example of the HFA (80-200 Hz) from Monkey 2. The x-axis is the time in the same range as a, and the y-axis is the electrode arrangement, which are sorted in the same order as a. **c.** Ratio of electrodes showing HFOs (HFA>3 SD) at each time point. Horizontal lines indicate the ratio of 0.5 criterion for cortical-wide HFO (cwHFO). The x-axis is time, in the same range as in a. Vertical bars at the bottom of the figures indicate the times at which the cwHFO occurred.

**Figure S5. ECoG signals of Monkey 3 on different days after the injection**.

ECoG signals of Monkey 3 on days 18 (left) and 38 (right) after the injection, similar to Figure S4. **a,** Examples of raw ECoG signals over a period of 2 min. **b,** Examples of HFA (80-200 Hz) of Monkey 3. **c,** Ratio of electrodes showing HFOs (HFA>3 SD) at each time point. The vertical bars at the bottom indicate the onset of the cwHFOs, as in the previous figure.

**Figure S6. cwHFOs following HFOs in V1 (left) and TC (right) in Monkey 3.** The cwHFOs followed the HFOs in the V1 and TC in Monkey 3. **a.** Left Column, 18 days after injection, **b.** Right Column, 38 days after injection. Top: Examples of HFA (80-200 Hz). The vertical bars at the bottom of the figures indicate the onset of the cwHFOs. The filled triangles correspond to the onset of the cwHFO magnified in the lower panels. Bottom: Examples of magnified cwHFO following an HFO in V1 (left) and TC (right) in a short time window (from -0.5 to 1 s).

**Figure S7. ECoG signals of Monkey 3 late (145 days) after KD. a,** Examples of raw ECoG signals over periods of 2 min (left) and 10 s (right). The shaded area on the x-axis of the left figure corresponds to the time range of the right figure, which are also shown in the following figures (b) and (c). **b,** Examples of HFA (80-200 Hz). The x-axis is time, corresponding to the same range as in a, and the y-axis is the electrodes, which are sorted in the same order as in a. **c,** Ratio of electrodes showing HFOs (HFA>3 SD) at each time point. The thin horizontal line indicates the 0.5 criterion for a cwHFO. The x-axis is time, corresponding to the same range as in a, and the vertical bars at the bottom of the figures indicate the onset of the cwHFOs.

**Figure S8. Analysis of glial and cell death markers in Monkey 4. a,** TOP left: Analysis of the glial and cell death markers in Monkey 4 at the *shCRTC1*#1 and *shscr* injection sites shown as filled blue circles and red circles, respectively. Top right: Nissl staining. In Monkey 4, we observed only local IEG expression (Supplemental Table 1a). IBA1 (middle row) and GFAP (bottom row) expression are shown in magenta. The merged image of either IBA1 or GFAP signals with Sirius signals is shown in the right column. **b,** Top left: the injection sites of *shCRTC1*#1 and *shscr*, left and right, respectively. GFAP (magenta) and Sirius (green). TUNEL signals were detected only at the *shCRTC1*#1 injection site. The square region on the right panel is magnified in the lower left view. **c,** Left: Merged signals with Sirius (green) and NeuN signals. Right: Only NeuN signals are shown. Layers 1, 2, 6 and white matter are abbreviated as L1, L2, L6, and wm, respectively.

**Figure S9. Expression of cell death related markers and IEGs at each shCRTC1 variant injection site. a,** Left: Injection sites of *shCRTC1*#1, #2, #3, #4, #5 (blue circles) and *shscr* (scrambled shRNA, red circles) in Monkey 6 are shown from a posterior view similar to that shown in Fig. 1b. Right: Experimental protocol. **b-g,** Histological analysis around the injection sites in V1 of Monkey 6. (b) IHC signals of Sirius (left) and CRTC1 (right). Adjacent sections were analyzed. Top: *shCRTC1*#5, middle: *shCRTC1*#1, *shscr* and *shCRTC1*#3, bottom: *shCRTC1*#2 and *shCRTC1*#4. Scale bars (1 mm). (c) Double IHC of NeuN (magenta) and Sirius (green). Top: Merged images of NeuN and Sirius signals. Bottom: NeuN signals. Left column: shCRTC1#3, middle column: shCRTC1#1, shscr and shCRTC1#5, right column: shCRTC1#4 and shCRTC#2, from left to right. Scale bars (1 mm). (d) Nissl stained sections. Top left row: *shCRTC1*#1, *shscr*, and *shCRTC*#5 (from left to right). Top right row: *shCRTC1* #4 and #2 (left and right, respectively). Middle row: enlarged *shCRTC1*#1, *shscr* and *shCRTC*#2 (from left to right). Bottom: *shCRTC1*#5 and #4 (left and right, respectively). Scale bars (1 mm). (e) Double IHC of glial markers and Sirius. Top two panels: GFAP (magenta), Sirius (green). Bottom two panels: IBA1 (magenta), Sirius (green). Nissl staining (d) was performed on the same sections, which were immunostained for IBA1 and Sirius. The IBA1 signal at the *shCRTC1*#5 injection site is due to the trace of the glass needle during injection. (f) Top: Expression of pCREB at each injection site for from *shCRTC1*#1 to *shCRTC1*#5. Bottom: Sirius. Adjacent sections were analyzed. Scale bar (500 μm). (g) Double ISH of *cFOS* (magenta) and *Sirius* (green). Top: merged images. The injection sites of *shCRTC1*#1 (left), *shscr* (middle), *shCRTC1*#3 (right). Bottom: *cFOS* expression (magenta); arrowheads indicate the borders of each injection. Note that sections of (b) and (g) were obtained from the L hemisphere and sections from (c) to (f) were obtained from the R hemisphere. Adjacent sections were used for (c) to (f). **h,** Top left: an illustration of the injection sites of Monkey 7 in the same style as in (a). Green dotted line of L hemisphere: the direction of sectioning. Top right: merged image of double IHC of Sirius and CRTC1 (injection sites: *shCRTC1*#3, *shscr* and #2, left, middle and right, respectively). Bottom: ISH of *ZIF268*, corresponding to the upper right section. The red arrow indicates the position of *shscr* injection site. Blue dotted line shows V1/V2 border. Scale bar (500 μm). **i,** Upper three panels (from left to right). Left: merged image of double IHC images of CRTC1 and Sirius (injection sites: *shCRTC1*#3, *shscr*, *shCRTC1* #5 and #2, from left to right). Middle: Backlit image. Right: IHC of GFAP. Bottom Left two panels: Enlarged IHC images of merged of IBA1 and Sirius (left), and IBA1 (right) at the *shCRTC1*#2 injection site. Bottom right two panels: GFAP (left). TUNEL staining (right): The *shCRTC1#2*-injected area is surrounded by the thin blue line. TUNEL signals are marked with small red dots within this blue line. White and black arrows in the upper panels indicate the same injection site of *shCRTC1#2*. Abbreviations: L1 (layer 1), WM (white matter). Scale bars (1 mm).

**Figure S10. A possible pathway for cell death and IEG induction by shCRTC1.** We sugest a possible pathway that explains how shCRTC1 KD causes cell death signaling. ① Synaptic activity causes Ca2+/calmodulin (CaM) dephosphorylation of CRTC1 mediated by calcineurin (Cn) and NF-AT41^1^. ② Dephosphorylated CRTC1 is transported to the nucleus^2,3,4,5^. ③ In the nucleus, CRTC1 activates both phosphorylated and dephosphorylated CREB^2^. ④ CRTC1 and the CREB complex activate SIK1 transcription, whose expression then activates the phosphorylation of CRTC1^2^. ⑤ The phosphorylation of CRTC1 leads to its export to the cytoplasm^2^. Note that there are two parallel pathways that activate the CREB-dependent cre promoter^2^ and thereby control IEGs, including cFOS and SIK1. ⑥ One is likely to depend on CaM kinase (CaMKK) and CaMKIV-dependent CREB phosphorylation^6^. The other is CRTC1-dependent CREB phosphorylation^2^ (③). Either phosphorylated CREB or nonphosphorylated CREB can activate cre dependent IEG expression together with CRTC1, both of which pathways are thought to be reduced by CRTC1 KD. ⑦ A few reports have implicated SIK1 in epilepsy^7,8,9^. SIK1 is a class II HDAC kinase and promotes skeletal myocyte survival^10^. ⑧ Upregulation of cFOS expression with neuronal apoptosis^11,12^ and induction of apoptosis by cFOS have been reported^13^. ⑨ Apoptosis/cell death and the induction of IEGs may cause self-amplification. ⑩ The abnormal expression of IEGs then causes abnormal activities that spread outside the cell.

**Supplemental Table 1**

a. **List of monkeys injected with 5 different shCRTC1**

Each row shows the marmosets used in this study. Each row indicates the experiment and animal characteristics. From left to right: (1) Monkey ID, (2) Sex (Male, Female), (3) Age (month), (4) Experimental period: injection and sacrifice between days, (5) Hemisphere, (6) IEG expression at shCRTC1s injection sites. Normal means a pattern similar to the non-injected hemisphere. Diffusible means that IEGs expression is widespread in the injected hemisphere, as shown in Figure 1d. Localized means that IEGs were expressed only at the site of injection, as shown in Figure 2. Reduced means that the IEGs expression was reduced only at the injection site, as shown in Figure 3. (7) Numerical values. The number of injection sites for each shRNA type (*shscr,* *shCRTC1* from #1 to #5) is shown below the column title for each of the two hemispheres. (8) Notes Experimental conditions for each monkey (see text for details).

b. **ISH probe list**

See the ISH section in STAR METHODS

**Supplemental Videos**

(1) **Supplemental Video 1**: ECoG recording of monkey 5 on day 22 after the injection

See the legend of Extened Data Fig 7b.

(2) **Supplemental Video 2**: ECoG recording of monkey 5 on day 39 after the injection

See the legend of Extened Data Fig 7b.

**Supplemental References**

1.	Shibasaki, F., Price, E.R., Milan, D., and McKeon, F. (1996). Role of kinases and the phosphatase calcineurin in the nuclear shuttling of transcription factor NF-AT4. Nature 382, 370-373. 10.1038/382370a0.

2.	Nonaka, M., Kim, R., Fukushima, H., Sasaki, K., Suzuki, K., Okamura, M., Ishii, Y., Kawashima, T., Kamijo, S., Takemoto-Kimura, S., et al. (2014). Region-specific activation of CRTC1-CREB signaling mediates long-term fear memory. Neuron 84, 92-106. 10.1016/j.neuron.2014.08.049.

3.	Uchida, S., Teubner, B.J.W., Hevi, C., Hara, K., Kobayashi, A., Dave, R.M., Shintaku, T., Jaikhan, P., Yamagata, H., Suzuki, T., et al. (2017). CRTC1 Nuclear Translocation Following Learning Modulates Memory Strength via Exchange of Chromatin Remodeling Complexes on the Fgf1 Gene. Cell Rep 18, 352-366. 10.1016/j.celrep.2016.12.052.

4.	Ch'ng, T.H., Uzgil, B., Lin, P., Avliyakulov, N.K., O'Dell, T.J., and Martin, K.C. (2012). Activity-dependent transport of the transcriptional coactivator CRTC1 from synapse to nucleus. Cell 150, 207-221. 10.1016/j.cell.2012.05.027.

5.	Jagannath, A., Butler, R., Godinho, S.I.H., Couch, Y., Brown, L.A., Vasudevan, S.R., Flanagan, K.C., Anthony, D., Churchill, G.C., Wood, M.J.A., et al. (2013). The CRTC1-SIK1 pathway regulates entrainment of the circadian clock. Cell 154, 1100-1111. 10.1016/j.cell.2013.08.004.

6.	Bito, H., Deisseroth, K., and Tsien, R.W. (1996). CREB phosphorylation and dephosphorylation: a Ca(2+)- and stimulus duration-dependent switch for hippocampal gene expression. Cell 87, 1203-1214. 10.1016/s0092

7.	Proschel, C., Hansen, J.N., Ali, A., Tuttle, E., Lacagnina, M., Buscaglia, G., Halterman, M.W., and Paciorkowski, A.R. (2017). Epilepsy-causing sequence variations in SIK1 disrupt synaptic activity response gene expression and affect neuronal morphology. Eur J Hum Genet 25, 216-221. 10.1038/ejhg.2016.145.

8.	Hansen, J., Snow, C., Tuttle, E., Ghoneim, D.H., Yang, C.S., Spencer, A., Gunter, S.A., Smyser, C.D., Gurnett, C.A., Shinawi, M., et al. (2015). De novo mutations in SIK1 cause a spectrum of developmental epilepsies. Am J Hum Genet 96, 682-690. 10.1016/j.ajhg.2015.02.013.

9.	Pang, B., Mori, T., Badawi, M., Zhou, M., Guo, Q., Suzuki-Kouyama, E., Yanagawa, T., Shirai, Y., and Tabuchi, K. (2022). An Epilepsy-Associated Mutation of Salt-Inducible Kinase 1 Increases the Susceptibility to Epileptic Seizures and Interferes with Adrenocorticotropic Hormone Therapy for Infantile Spasms in Mice. Int J Mol Sci 23. 10.3390/ijms23147927.

10.	Berdeaux, R., Goebel, N., Banaszynski, L., Takemori, H., Wandless, T., Shelton, G.D., and Montminy, M. (2007). SIK1 is a class II HDAC kinase that promotes survival of skeletal myocytes. Nat Med 13, 597-603. 10.1038/nm1573.

11.	Kalra, N., and Kumar, V. (2004). c-Fos is a mediator of the c-myc-induced apoptotic signaling in serum-deprived hepatoma cells via the p38 mitogen-activated protein kinase pathway. J Biol Chem 279, 25313-25319. 10.1074/jbc.M400932200.

12.	Chen, X., Shen, J., Wang, Y., Chen, X., Yu, S., Shi, H., and Huo, K. (2015). Up-regulation of c-Fos associated with neuronal apoptosis following intracerebral hemorrhage. Cell Mol Neurobiol 35, 363-376. 10.1007/s10571-014-0132-z.

13.	Preston, G.A., Lyon, T.T., Yin, Y., Lang, J.E., Solomon, G., Annab, L., Srinivasan, D.G., Alcorta, D.A., and Barrett, J.C. (1996). Induction of apoptosis by c-Fos protein. Mol Cell Biol 16, 211-218. 10.1128/MCB.16.1.211.
