## Supplementary figures and images for "Epileptic responses in marmosets induced by knockdown of CREB Regulated Transcription Coactivator 1 (CRTC1)"

### Figure S1

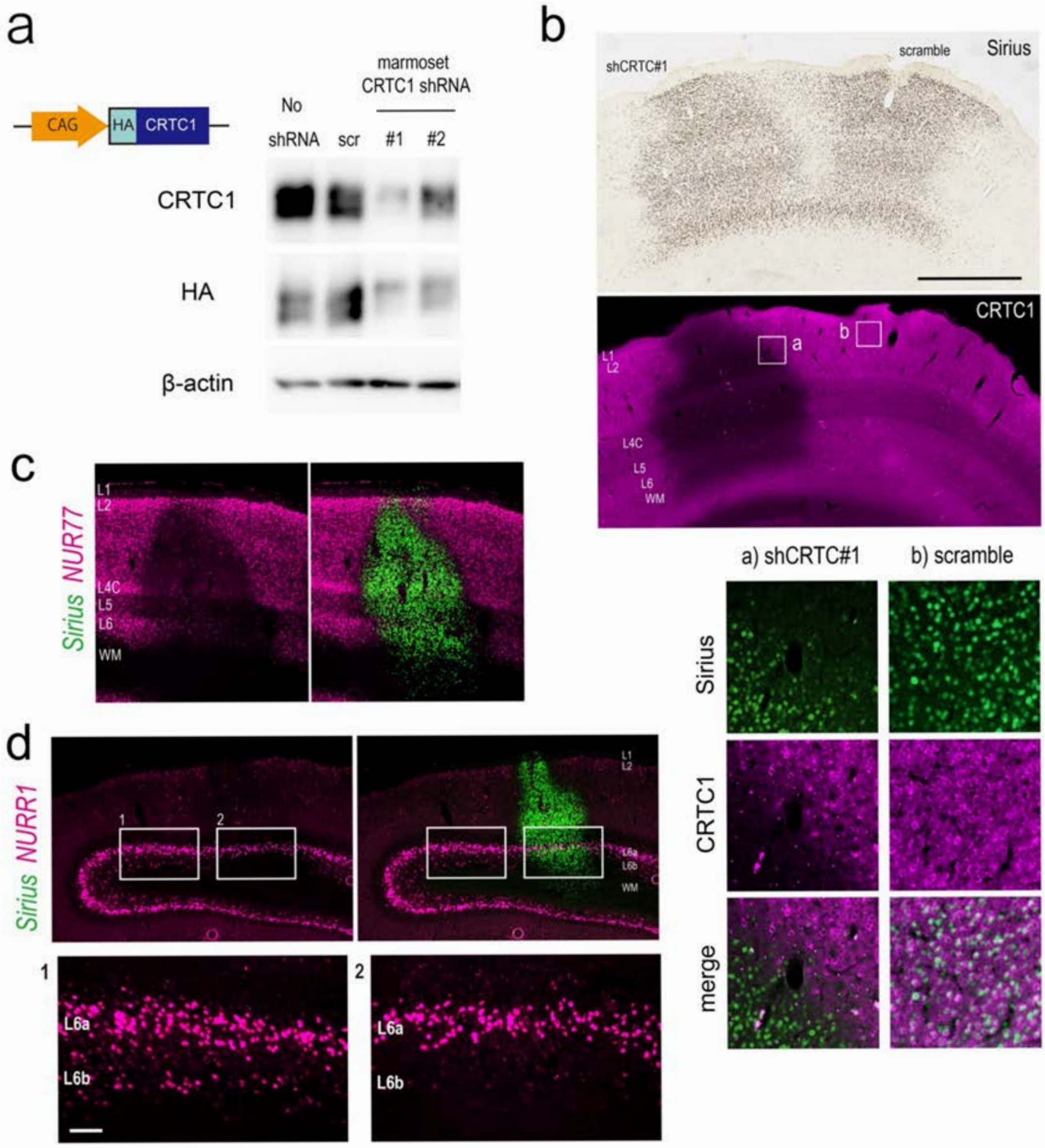

### Figure S2

*Figure S2*

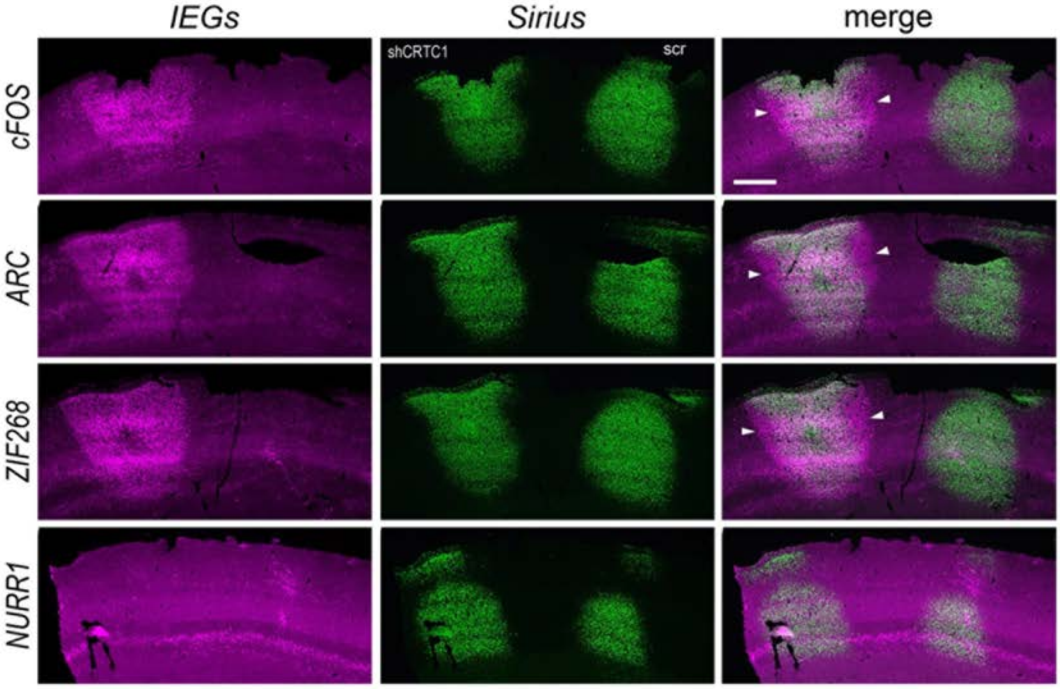

### Figure S3

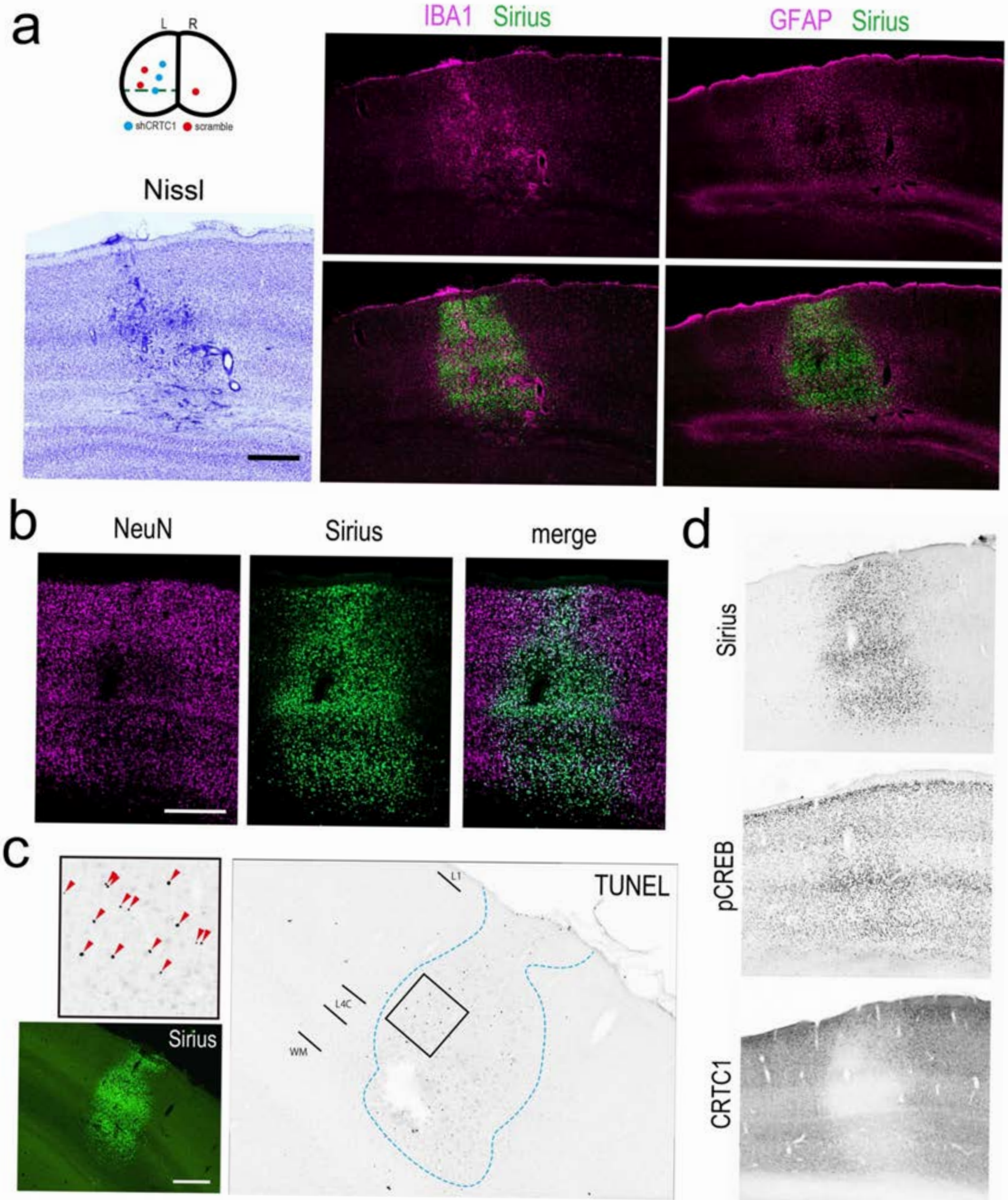

### Figure S4

*Figure S4*

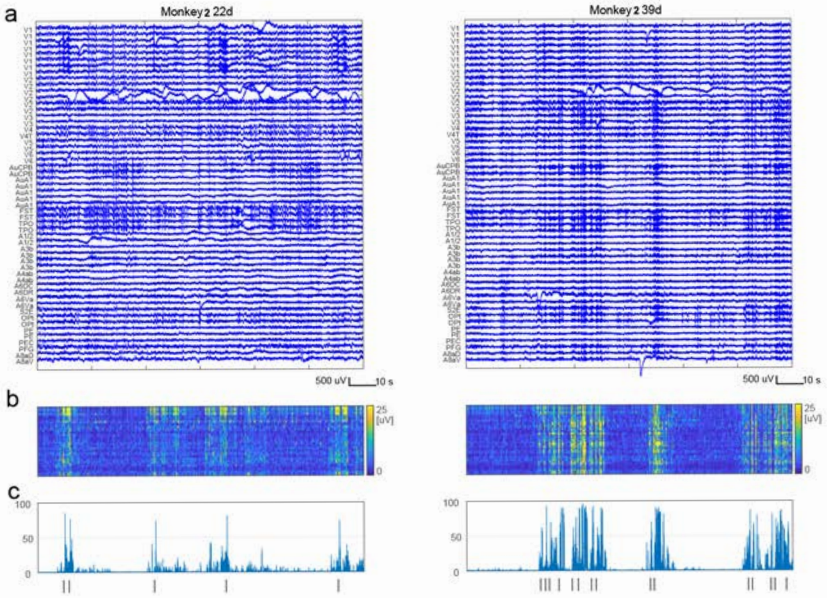

### Figure S5

Figure S5

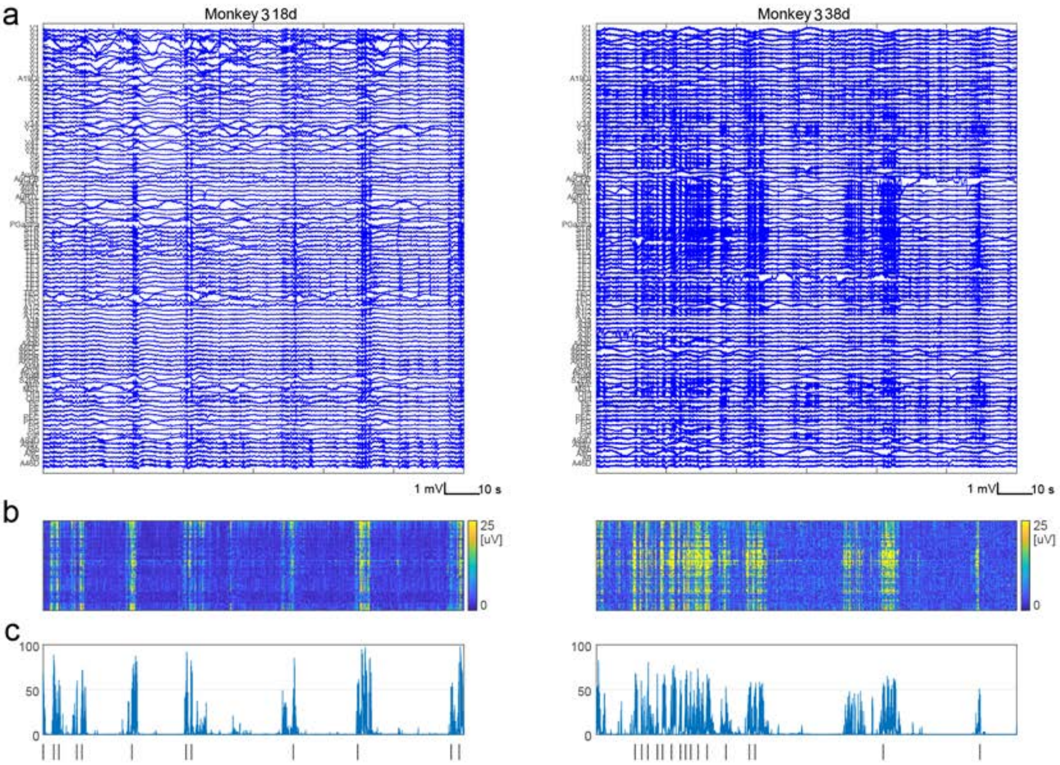

### Figure S6

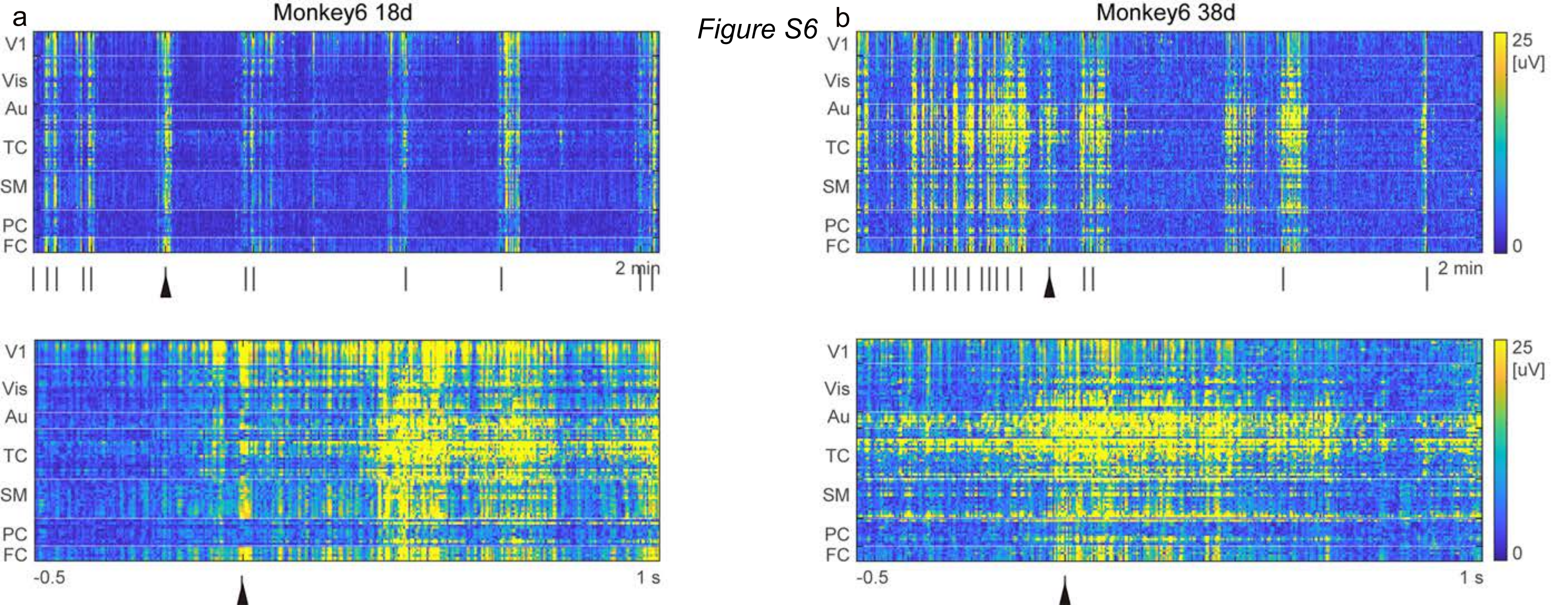

### Figure S7

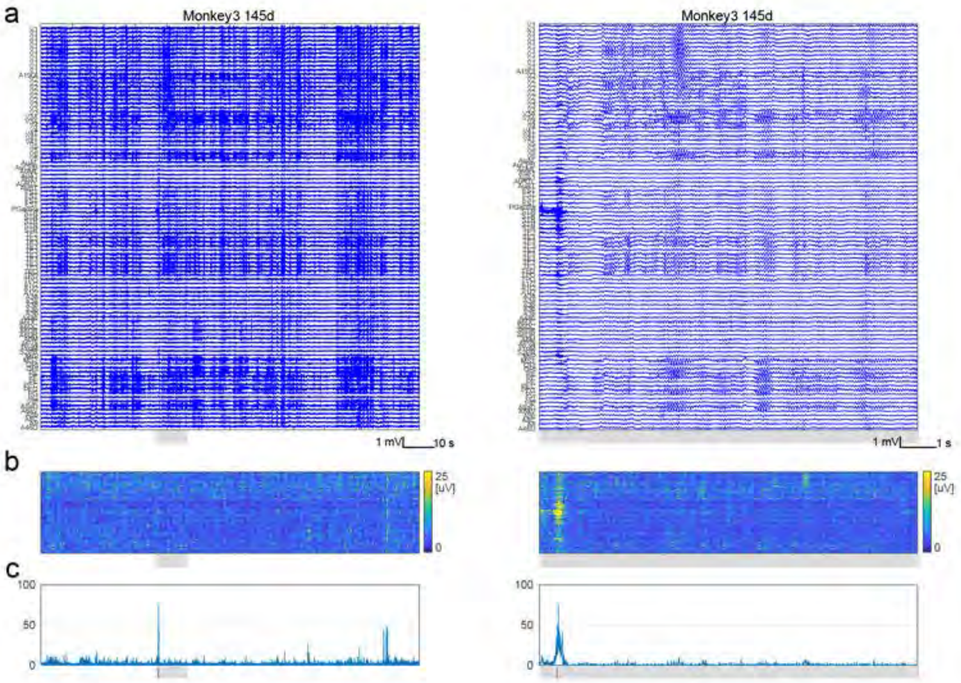

### Figure S8

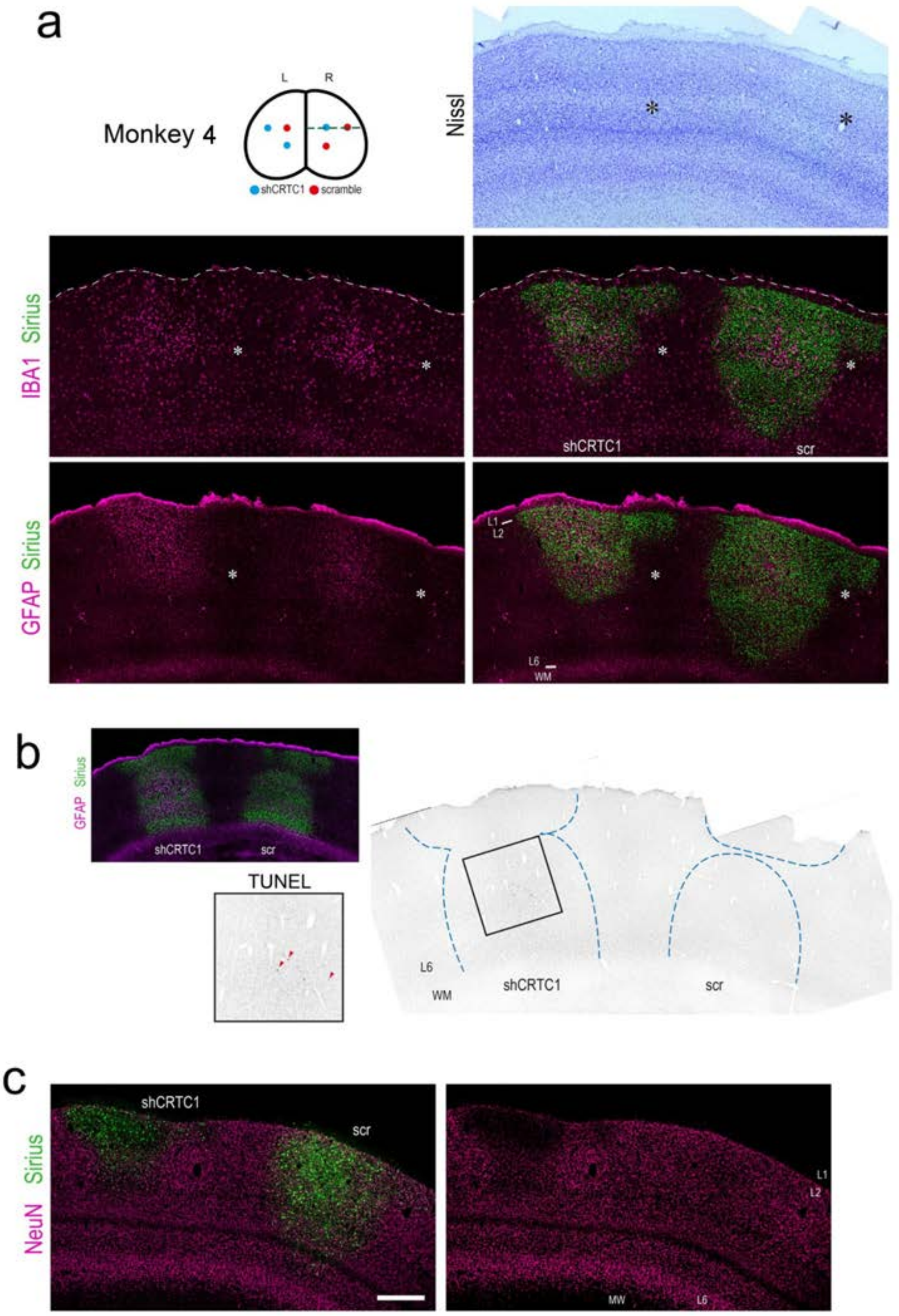

### Figure S9

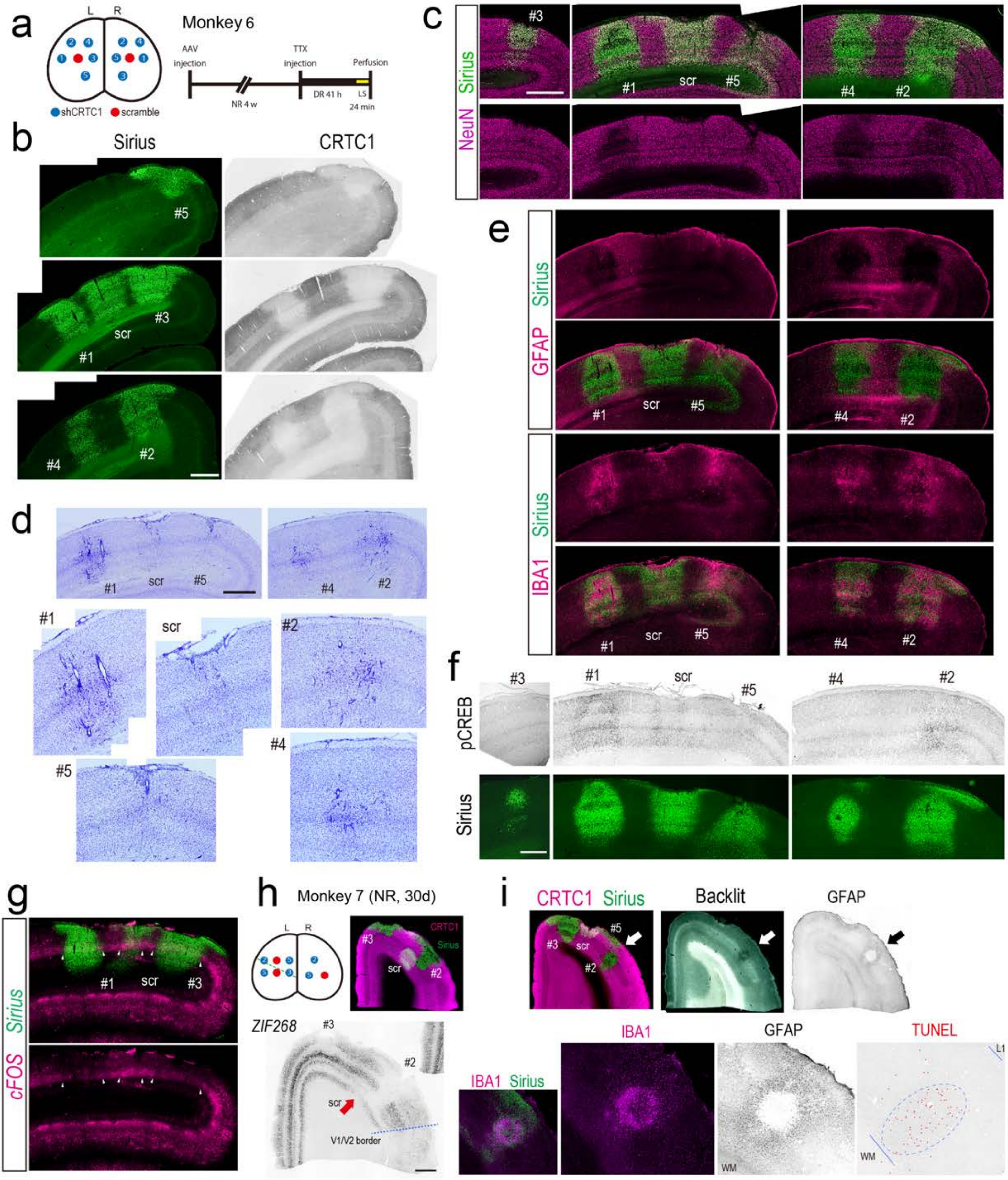

### Figure S10

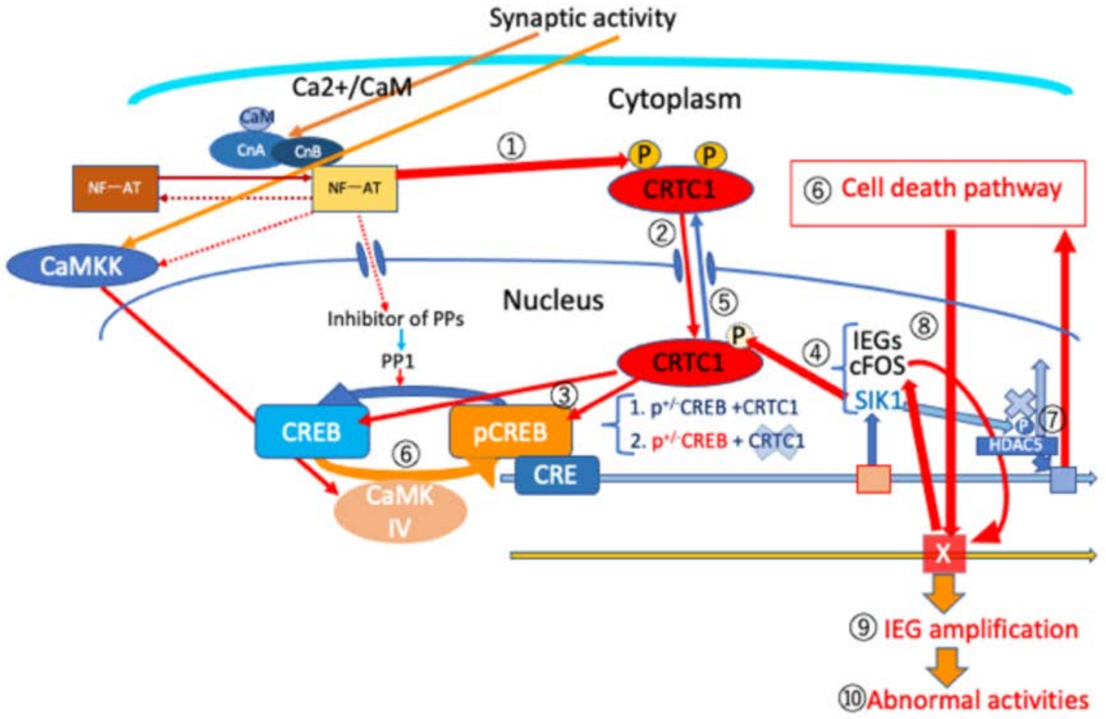

### Supplemental Video1, 2.pptx

#### Slide 1
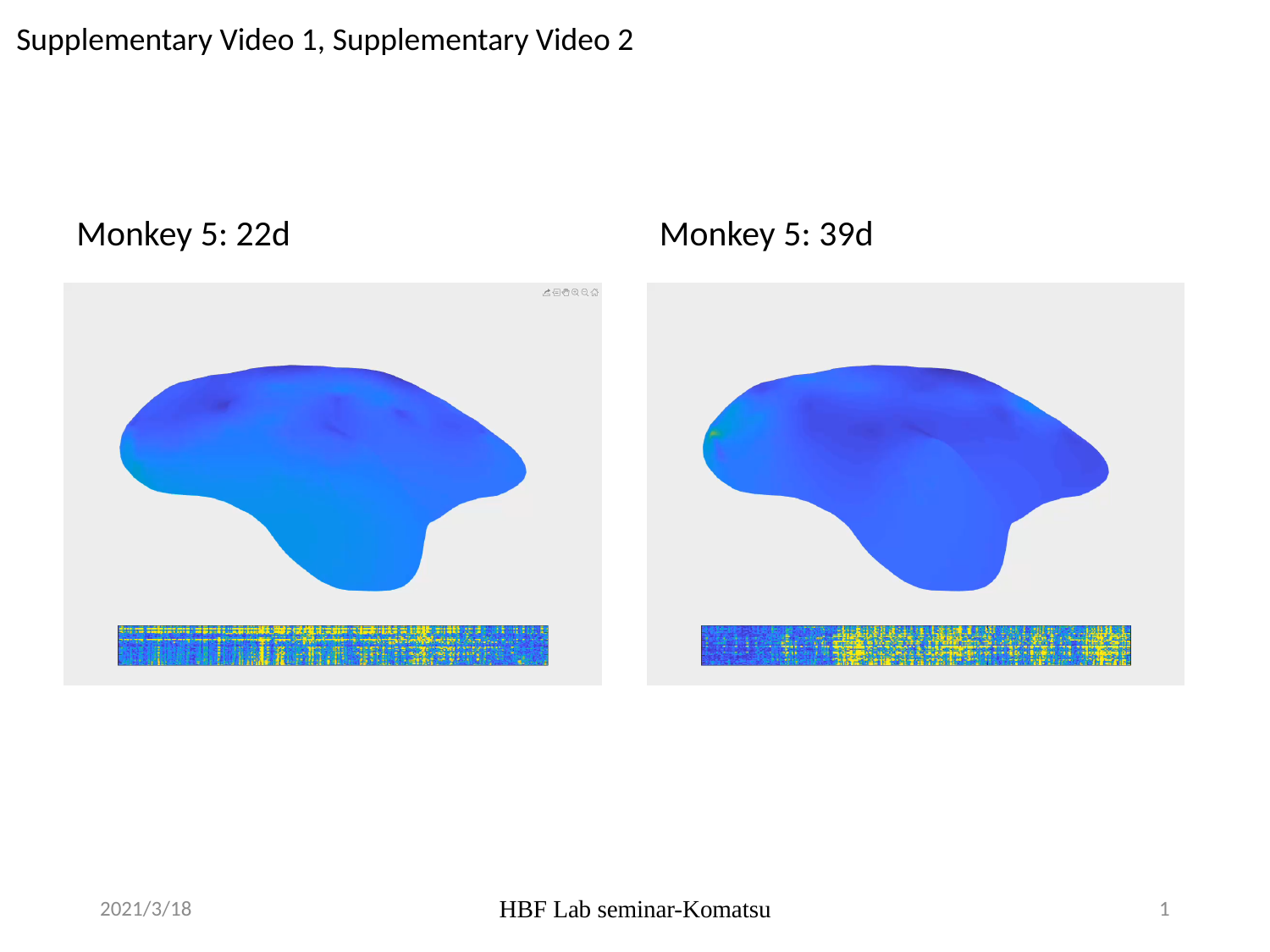

Supplementary Video 1, Supplementary Video 2
Monkey 5: 22d
Monkey 5: 39d
2021/3/18
HBF Lab seminar-Komatsu
1
